# Genomics of the diversification of *Heliconius cydno* (Lepidoptera: Nymphalidae) in northern South America: geographic and phenotypic patterns

**DOI:** 10.64898/2026.11.20.768178

**Authors:** Alexandra Arias-Cárdenas, Nicol Rueda-M, Carlos F. Arias, Mark Ravinet, Owen McMillan, Carolina Pardo-Diaz, Camilo Salazar

**Affiliations:** School of Sciences and Engineering, Biology Program, Universidad del Rosario, Bogotá D.C., Colombia; Tree of Life Programme, Wellcome Sanger Institute, Hinxton, United Kingdom; Smithsonian Tropical Research Institute, Panama City, Panamá; Centre for Ecological and Evolutionary Synthesis, Department of Biosciences, University of Oslo, Oslo 0316, Norway

**Keywords:** Adaptive radiation, *Heliconius cydno*, phylogeography, historical demography, population genomics, *cis*-regulatory modular architecture, white submarginal band

## Abstract

The interplay between natural selection, gene flow, and geography is a key driver of population differentiation. This study investigates their influence on the recent adaptive radiation of *Heliconius cydno*, a mimetic and aposematic butterfly with striking variation in wing color pattern across its distribution in Central and northern South America. The genetic basis of this phenotypic diversity has a modular architecture, whereby a few key genes and *cis*-regulatory elements control differences among morphotypes. Using whole-genome sequences from 133 individuals representing all subspecies of *H. cydno*, we investigated (1) the role of Andean orogeny in its diversification and (2) the genetic basis of the variation in the hindwing white submarginal band, which to date, had remained unknown. First, we found that five of the fourteen described subspecies correspond to distinct monophyletic lineages and well-defined genetic clusters that originated in northwestern South America and later expanded into Central America and the Colombian inter-Andean valleys. Second, we observed signatures of genetic structure that were consistent with geographic patterns, particularly those associated with the inter-Andean region and Gorgona Island. Third, we identified *ivory:mir-193* as the locus underlying variation in the white submarginal band on the hindwing. We also detected evidence of introgression suggesting that this trait was likely transferred from *H. cydno* to *H. melpomene*, contributing to the establishment of a novel mimicry ring with *H. erato*. Altogether, these findings highlight the roles of geography, genomic architecture, and introgression in the diversification of *H. cydno*.

## INTRODUCTION

Recent adaptive radiations, characterized by rapid diversification and the evolution of contrasting phenotypes suited to different environments (Gavrilets & Vose, 2005; Losos, 2010; Berner & Salzburger, 2015; Marques *et al*., 2018), offer a powerful framework to study how evolutionary forces, local adaptation, and demographic processes interact during the speciation process. Yet, the spatial and temporal patterns and mechanisms of diversification vary widely among lineages (Gillespie *et al*., 2020), underscoring the need to study early diversification within species to understand what drives phenotypic diversity and speciation.

Radiations range from intraspecific divergence to ancient splits among major lineages (Gillespie *et al*., 2020), shaped by a mix of adaptive and non-adaptive mechanisms (Matsubayashi & Yamaguchi, 2022). Many documented cases involve high speciation rates linked to substantial genomic divergence, chromosomal inversions, or introgression, such as Anolis lizards in the Antilles (Losos, 2011; Tollis *et al*., 2018), Darwin’s finches in the Galápagos Islands (Almén, 2016), and cichlids from Lake Victoria (Blumer *et al.,* 2025; Marques *et al*., 2025). In other cases, adaptive processes and/or sexual selection drive diversification among recently diverged species with low genomic divergence, as in hamlet fishes (<10,000 generations; Hench *et al.,* 2022) and Galápagos tree finches (1–2 Ma; Lamichhaney *et al*., 2015), where color pattern and beak differences, respectively, map to a few genomic regions against an otherwise undifferentiated genomic background (Helmkampf *et al*., 2025; Sebastianelli-Arbelaez *et al*., 2026). Radiation can even occur within species, among low-divergence populations or morphs lacking reproductive isolation, without leading to speciation (Calsbeek *et al*., 2007; Levis *et al*., 2017).

Although adaptive radiations are best documented in insular or lacustrine systems (Cerca *et al*., 2023), they also occur in continental settings with high geographic and ecological complexity, such as the Neotropics, a region that hosts major adaptive radiations across riverine cichlid fishes, rodents, ants, *Espeletia* plants, Ithomiinae and *Heliconius* butterflies (Guevara-Andino *et al*., 2025; López-Fernández *et al*., 2013; Price *et al*., 2014; Pouchon *et al*., 2018; Cicconardi *et al*., 2023; Vallejos-Garrido *et al*., 2023; Van der Heijden *et al*., 2025). Yet continental radiations in this megadiverse region remain relatively understudied (Raven *et al*., 2020). In the Neotropics, the gradual and discontinuous uplift of the Andes reshaped South American landscapes and promoted speciation (Chapman, 1917; Hoorn *et al*., 2010), as orogeny-induced vicariance, dispersal corridors, and environmental heterogeneity constrained gene flow, favored local adaptation, and drove genetic differentiation (Pelletier & Carstens, 2018; Jensen *et al*., 2024), positioning the Neotropics as an ideal system to investigate the processes driving early-stage divergence in continental radiations. However, studies of low-divergence neotropical species have focused mainly on abiotic and geographic drivers of diversification (Rojas *et al*., 2019; Salgado-Roa *et al*., 2024; Sánchez *et al*., 2024), leaving the genetic basis of early divergence understudied even though elucidating the genetic architecture of traits involved in rapid adaptation is essential to understand how adaptive variation is maintained and shifts under selection (Whiting *et al*., 2022).

Diversification in Lepidoptera shows pronounced spatial and temporal heterogeneity, with the Andean region standing out as a center of origin and a major driver of diversification (Elias *et al*., 2009; Rosser *et al*., 2014; De-Silva *et al*., 2016, 2017; Chazot *et al*., 2016, 2018, 2019; Van der Heijden *et al*., 2025). In this region, the genus *Heliconius*, toxic and aposematic butterflies with extensive wing color pattern diversity and Müllerian mimicry, exemplifies this heterogeneity, illustrating how diversification depends on geographic patterns, demographic connectivity (Mallet & Gilbert, 1995; Mallet, 2000; Brower, 2013; Jiggins, 2017), and natural selection acting on a few large-effect loci underlying wing phenotypes (e.g. *optix*, *ivory*, *WntA* and *aristaless*; Martin *et al*., 2013; Westerman *et al*., 2018; Lewis *et al*., 2019; Livraghi *et al*., 2021, 2024). In these butterflies, diversification trajectories range from strongly structured populations shaped by reduced gene flow and geographic barriers to rapid expansions with persistent connectivity and weak population differentiation, partly reflecting variation in dispersal ability (Quek *et al*., 2010; Davey, 2013; Martin *et al*., 2016; Rodríguez de Cara *et al*., 2023; Seixas *et al*., 2024). These contrasting patterns are further shaped by frequency-dependent selection and assortative mating within mimicry systems (Jiggins *et al*., 2001; Kronforst et al., 2006; Chouteau *et al*., 2016, 2017; Moest *et al*., 2020; Rossi *et al*., 2024). In addition, adaptive color pattern loci and their *cis*-regulatory regions are recurrently shared across lineages through introgression, driving both phenotypic diversification and mimetic convergence (Pardo-Diaz *et al*., 2012; Dasmahapatra *et al*., 2012; Merrill *et al*., 2015; Wallbank *et al*., 2016; Enciso-Romero *et al*., 2017; Thawornwattana *et al*., 2023).

Studying diversification within a single *Heliconius* species allows us to investigate incipient divergence, where subspecies show adaptive phenotypic differences that may lead to reproductive isolation. Here, we focus on the recent South American radiation of *Heliconius cydno* (∼700 Kya; Guan *et al*., in prep.) to examine the evolutionary and demographic processes shaping its diversification and genetic structure, as well as the genomic basis of its phenotypic diversity. This species is distributed from Mexico to northern South America (Linares, 1996; 1997) and exhibits pronounced geographic variation in wing color patterns, particularly in the inter-Andean valleys of Colombia (Linares, 1996; 1997; Constantino & Salazar, 1998). This variation, linked to Müllerian mimicry, combines white or yellow forewing bands with either a white submarginal band or a yellow hindwing bar on an iridescent blue-black background (Linares, 1996; 1997). The species forms mimicry rings mainly with *Heliconius sapho* and *Heliconius eleuchia*, with phenotypic variation that is geographically restricted and only partially overlapping with that of its co-mimics, but it also forms local mimicry associations with other *Heliconius* species and with Lepidoptera from other genera (Linares, 1996, 1997).

Phylogeographic analyses based on mitochondrial (COI) and AFLP markers support the monophyly of *H. cydno* and identify three clades largely concordant with geography, though both markers show limited resolution within clades and low differentiation among subspecies (Arias *et al*., 2014). Also, assortative mating based on forewing color pattern has been documented: yellow males of the polymorphic race *H. c. alithea* preferentially court yellow over white females (Kronforst *et al*., 2006; VanKuren *et al*., 2025). Moreover, two races associated with distinct mimicry rings also show considerable genetic differentiation despite meeting in a unimodal hybrid zone (Linares, 1997; Merchán *et al*., 2005; Arias *et al*., 2012), where mating experiments have revealed incipient, asymmetric assortative mating driven by wing color pattern (Arias *et al*., 2012). Yet, despite extensive research on the genomic architecture of color pattern, the causal locus underlying the presence or absence of the white submarginal hindwing band of *H. cydno* remains unidentified (Livraghi *et al*., 2021; Hanly *et al*., 2026). Interestingly, across the distributions of *H. melpomene* and *H. erato* co-mimics, the white submarginal hindwing band is present only in *H. m. cythera* and *H. erato cyrbia*. This provides an opportunity to test whether the trait was acquired through introgression from *H. cydno*, where it is common, or originated de novo in *H. melpomene*. Either scenario could have contributed to the formation of a new mimetic ring.

Using whole-genome data, we characterize the adaptive radiation of *Heliconius cydno* through two approaches. First, we evaluate the role of geographic drivers (i.e., the Andes and inter-Andean valleys) in the origin of the radiation and the genetic structure of the species and infer the demographic processes shaping its diversification. Second, we explore the genetic architecture of its phenotypic diversity, focusing on the locus underlying the white submarginal hindwing band and testing for introgression with *H. melpomene*. Together, these analyses contribute to understanding how landscape and genetic variation shape adaptive phenotypes during the early stages of adaptive radiations.

## METHODS

### Sampling and whole-genome resequencing

Our sample set consisted of 133 *H. cydno* individuals spanning all described subspecies (Lamas *et al*., 2004), and an undescribed population from Gorgona Island, covering the full geographic range of the species. Of these, 79 individuals were newly sequenced for this study (Supplementary Fig. 1A; Supplementary Table 1), and 54 were obtained from published datasets. For the newly generated sequences, specimens were assigned to subspecies based on wing color pattern and geographic origin, following the taxonomic classification of Lamas & Jiggins (2016).

Genomic DNA was extracted from thoracic tissue using the DNeasy Blood & Tissue Kit (QIAGEN). DNA quality and integrity were assessed by fluorometry (Qubit, Thermo Fisher) and 1% agarose gel electrophoresis. Libraries were prepared and sequenced on the Illumina HiSeq X platform (Novogene, Beijing, China) at ∼30× coverage per individual.

### Read processing and genotype calling

Raw sequence quality was assessed using FastQC (Andrews, 2010) and paired-end adapters were removed using cutadapt v1.8.1 (Martin, 2011). Reads were mapped to the *H. melpomene* reference genome v2.5 available in LepBase (Challi *et al*., 2016), using BWA v0.7.17 with the BWA-MEM algorithm (Li, 2013). Aligned reads were indexed using SAMtools v1.8 (Li *et al*., 2009), and duplicate reads were removed with Picard v2.18.15 (Broad Institute, 2014).

Variants were called per individual with the GATK v4.1.9.0 HaplotypeCaller (DePristo *et al*., 2011) in BP resolution mode, followed by joint genotyping with GenotypeGVCFs. VCFtools v0.1.14 (Danecek *et al*., 2011) was used to compute summary statistics (allele frequency per site, mean depth per individual and per site, site quality, proportion of missing data, and mapping rate), and to retain sites with quality score ≥ 20 and <5% missing data. To remove likely paralogous regions, we excluded sites with sequencing depth <5 or mean depth per individual >1.5× the genome-wide average with the script *removeTooHighDepthSites*.*sh* (https://github.com/joanam/VictoriaRegionSuperflock/), and sites with excess heterozygosity across individuals with VCFtools v0.1.14 “*--hardy”* option and p < 1e−5 (Danecek *et al*., 2011).

The filtered VCF was split into 21 chromosomes using BCFtools v1.8 (Danecek *et al*., 2021), and haplotypes were phased per chromosome with SHAPEIT4 (Delaneau *et al*., 2019) using default parameters and chromosome-specific recombination rates (Davey *et al*., 2017).

### Subspecies tree and mtDNA network

We inferred the *H. cydno* subspecies tree using the coalescent-based method ASTRAL (Accurate Species TRee ALgorithm; Mirarab *et al*., 2014), which estimates a statistically consistent species tree from multiple gene trees. Gene trees were built from 2 Kb loci extracted from each chromosome, spaced ≥10 Kb apart to minimize linkage disequilibrium (Dasmahapatra *et al*., 2012), using a VCF file retaining monomorphic and polymorphic sites with VCFtools v0.1.14 (Danecek *et al*., 2011). We then generated 248 multilocus blocks of 100 concatenated loci using SAMtools v1.8 (Li *et al*., 2009), which were converted to PHYLIP format using vcf2phylip.py (https://github.com/joanam/scripts/blob/master/vcf2phylip.py) and used as input for IQ-TREE v2.2.0 (Nguyen *et al*., 2014) to infer maximum likelihood (ML) gene trees. The best-fitting nucleotide substitution model for each block was selected using ModelFinder (Kalyaanamoorthy *et al*., 2017), and node support was assessed with 1,000 ultrafast bootstrap replicates. The 248 resulting gene trees were used as input for ASTRAL to estimate the maximum quartet support subspecies tree. The analysis included all *H. cydno* subspecies, as well as two additional *Heliconius* species as outgroups: *H. melpomene* (subspecies *H. m. melpomene*, *H. m. malleti, and H. m. flavomartina*) and *H. timareta* (subspecies *H. t. florencia* and *H. t. thelxinoe*). Outgroups were mapped to the same reference genome and processed as described above.

The mtDNA scaffold was extracted from the VCF file using BCFtools v1.8 (Danecek *et al*., 2021), converted to FASTA format, and used to infer a mitochondrial haplotype network in SplitsTree4 v4.19.2 (Huson & Bryant, 2006), using Kimura two-parameter (K2P) genetic distances and the Neighbor-Net algorithm.

### Genetic structure, diversity statistics and ancestry analyses

Population structure among subspecies of *H. cydno* was assessed using multiple genome-wide approaches, excluding mtDNA. First, we conducted a Principal Component Analysis (PCA) using PLINK v2.0 (Chang *et al*., 2015) after removing sites in linkage disequilibrium (r² > 0.2 within 50 Kb sliding windows with a 10 Kb step size) and sites that were monomorphic, multiallelic, or had a low minor allele frequency (MAF < 0.02) with VCFtools v0.1.14 (Danecek *et al*., 2011).

Second, genetic structure and ancestry were assessed using maximum likelihood (ML) and Bayesian inference (BI). For the ML analysis, ADMIXTURE v1.3.0 (Alexander *et al*., 2009) tested K = 2 to 14, with the optimal number of clusters selected by the Akaike Information Criterion (AIC). For the BI analysis, FastSTRUCTURE (Raj *et al*., 2014) tested the same K range, with the most plausible K inferred using the *chooseK.py* script. To resolve population structure at a finer resolution, we used the haplotype-based method FineSTRUCTURE (Lawson *et al*., 2012), which estimates relationships among individuals from a coancestry matrix and infers each individual’s posterior probability of assignment to genetic clusters.

Third, we used Pixy v1.2.7.beta1 (Korunes & Samuk, 2021) to estimate nucleotide diversity per subspecies (π) and genetic differentiation between pairs of subspecies (relative differentiation F_ST_ and absolute differentiation Dxy). These genome-wide analyses were conducted in non-overlapping 50 Kb windows, including monomorphic and polymorphic sites. Absolute differentiation corrected for ancestral diversity (Da) was calculated from Dxy and π using custom scripts, and scatterplots of π versus Dxy and F_ST_ versus Dxy were used to compare subspecies across differentiation levels.

### Demographic analyses

We reconstructed the historical demography of this adaptive radiation using four complementary approaches. First, the species’ geographic origin was inferred with *X-ORIGIN* (He *et al*., 2017), which uses coalescent-based expansion simulations to estimate the probability that given geographic coordinates correspond to the origin of a population expansion, based on the spatial distribution of genetic diversity and the directionality index (Ψ; Peter & Slatkin, 2013). Default parameters were used, and results were visualized using RStudio v2024.09.1.

Second, historical migration patterns and potential barriers to gene flow were evaluated with Estimated Effective Migration Surfaces (EEMS; Petkova *et al*., 2016). Pairwise genetic dissimilarities were calculated using the *bed2diffs* script (https://github.com/dipetkov/eems), and EEMS was run with 200 demes across a polygon encompassing all sampling locations, using an MCMC of 10,000,000 iterations, discarding the first 10% as burn-in and sampling every 1,000 iterations.

Third, changes in effective population size over time were inferred using Relate v1.1.2 (Speidel *et al*., 2019), which estimates coalescence times from genealogies reconstructed from phased haplotypes using an ML framework. We used SHAPEIT4 (Delaneau *et al*., 2019) to generate per-chromosome input files (.haps/.sample) that were transformed to Relate format using *PrepareInputFiles.sh.* The .anc and .mut files were generated using Relate’s “--mode ALL” option with an effective population size of 2×10⁷ and a mutation rate of 2.9×10⁻⁹ per site per generation from *H. melpomene* (Keightley *et al*., 2015). Effective population size trajectories were inferred with *EstimatePopulationSize.sh*, using a previously estimated genetic map for *H. cydno* (Martin *et al*., 2019).

Finally, Tajima’s D was calculated genome-wide per subspecies using VCFtools v0.1.14 (Danecek *et al*., 2011), in non-overlapping 50 Kb windows including monomorphic and polymorphic sites, to detect genomic signatures of demographic changes, such as population expansions or bottlenecks.

### Genetic basis of the white submarginal band

As a preliminary step, we conducted a genome-wide F_ST_ scan between the subspecies pair with the greatest wing pattern differences (*H. c. cydnides* and *H. c. weymeri*; Supplementary Fig. 1B) to confirm that our sampling design had sufficient power to recover known divergence peaks (including major color pattern loci). We then used three complementary approaches to identify genomic regions associated with the white submarginal band.

First, genome-wide F_ST_ was estimated with Pixy v1.2.7.beta1 (Korunes & Samuk, 2021) in non-overlapping 50 Kb windows (monomorphic and polymorphic sites), grouping *H. cydno* subspecies by presence or absence of the white submarginal band (Supplementary Fig. 1B; Supplementary Table 1) and excluding *H. c. subsp. nov.* due to its strong genetic divergence. Because linkage between the *Yb* and *Sb* loci (associated with the yellow hindwing bar and the white submarginal band, respectively) has been reported in *H. cydno* and *H. cydno* × *H. melpomene* broods (Linares, 1997; Naisbit *et al*., 2003; Merchán *et al*., 2005), and regulatory elements for the yellow hindwing bar map to chromosome 15 (Enciso-Romero *et al*., 2017; Livraghi *et al*., 2024), subsequent F_ST_ analyses focused on this chromosome. To control for population structure on this chromosome, F_ST_ scans comparing the presence versus absence of the white submarginal band were run within each of the following geographic regions: Central America, Pacific, and Cauca Valley. All four subspecies from the Magdalena Valley (*H. c. lisethae, H. c. cydno, H. c. wanningeri, H. c. hermogenes*) have the white submarginal band and were compared, as a group, with the Venezuelan subspecies lacking the band (*H. c. cordula, H. c. gadouae*) using 2 Kb windows; *H. c. barinasensis* was excluded due to low sample size (n = 2). We estimated Dxy, π (Pixy v1.2.7.beta1; Korunes & Samuk, 2021), and Tajima’s D (VCFtools v0.1.14; Danecek *et al.,* 2011) at the same window size, and their statistical significance was assessed via permutation tests with the *permTests* function in the R package *regioneR* (Gel *et al*., 2016) against 50,000 randomly sampled 2 Kb regions genome-wide, comparing weighted means in the candidate region with genome-wide expectations.

Second, a genome-wide association analysis (GWAS; Uffelmann *et al*., 2021) was run with GEMMA v0.98.5 (Zhou & Stephens, 2012) using a univariate linear mixed model with a pairwise relatedness matrix as a covariate. This was done first genome-wide and then restricted to chromosome 15, grouping individuals by phenotype and geography; significance was assessed with a Bonferroni correction (0.05/number of SNPs). Spearman’s rank correlation coefficient (ρ²) was calculated for significant SNPs using custom scripts, and the most strongly associated SNPs were visualized in a genotype matrix with the R package *GenotypePlot v0.2.1* (Whiting, 2022) to confirm their association with the presence or absence of the white submarginal band.

Third, we applied a topology-weighting approach with Twisst (Martin & Van Belleghem, 2017) to the candidate region associated with the white submarginal band. We tested three hypotheses: (i) clustering by phenotype (band presence/absence), (ii) clustering by geography, and (iii) a random topology, across four pairwise comparisons: (i) Central America vs. Pacific, (ii) Pacific vs. Cauca Valley, (iii) Central America vs. Cauca Valley, and (iv) Pacific vs. Magdalena Valley + Venezuela. Maximum likelihood trees were inferred with RAxML v8.2.9 (Stamatakis *et al*., 2008) in sliding windows of 50 SNPs (*raxml_sliding_windows.py*), retaining windows with ≥50% genotyped individuals. Twisst was run with “-*-method complete*” and default parameters.

We also tested for signals of selection by estimating the composite likelihood ratio (CLR) with SweepFinder2 (SF2; DeGiorgio *et al*., 2016), using VCF files with monomorphic and polymorphic sites and *H. m. melpomene* as the outgroup, and grouping subspecies by geographic region (described above). SF2 was run in two modes per dataset: with the region-specific site frequency spectrum (SFS; inferred with the -sg option) and with a genome-wide SFS excluding the candidate region as a neutral null model (-lg option), using the same 2 Kb window size as above. CLRs and 1/α values were visualized in RStudio v2024.09.1 with custom scripts.

Because the mimetic hindwing color pattern loci are in repulsion in *H. cydno* subspecies (Supplementary Fig. 1B), we estimated LD between the white submarginal band candidate region and previously identified cis-regulatory elements (CREs) controlling the hindwing yellow bar (Enciso-Romero *et al*., 2017; Livraghi *et al*., 2024). A pairwise SNP correlation matrix (r²) was obtained with PLINK v2.0 (Chang *et al*., 2015), retaining SNP pairs <1 Mb apart and reporting those with r² ≥ 0.2. Loci positions were defined as *Yb(d)*: 1,403,500–1,405,500 and *Yb(v)*: 1,429,500–1,430,500 (Livraghi *et al*., 2024), and the *Sb* regulatory region as estimated in this study. Analyses included individuals of *H. c. weymeri* (yellow bar) and *H. c. cydnides* (white submarginal band), which co-occur geographically (Fig. 1A).

**Figure 1.**
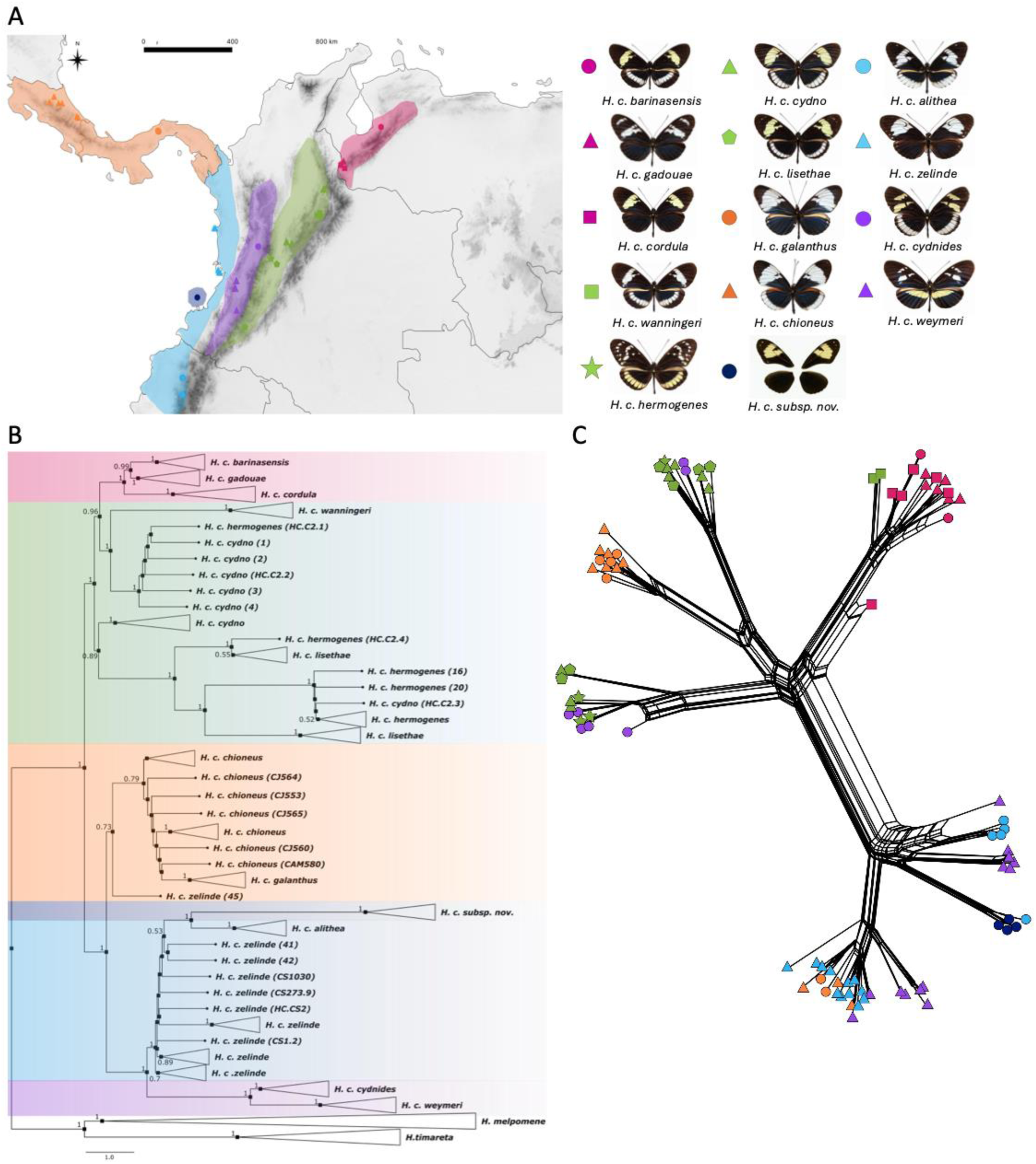
Geographic distribution, phenotypic variation, and phylogenetic relationships among *H. cydno* subspecies. **A)** Geographic distribution of subspecies and their phenotypes, grouped according to the geographic regions identified in the phylogenetic analyses: Central America (orange), Pacific (light blue), Cauca Valley (purple), Magdalena Valley (green), Venezuela (pink), and Gorgona (dark blue). **B)** ASTRAL subspecies tree inferred from 248 trees. Clades are colored according to the geographic regions shown in panel A. Nodes indicate posterior probability values as measures of support, and branch lengths correspond to coalescent units. **C)** SplitsTree4 network based on mtDNA, with tips represented by geometric shapes according to subspecies and colored according to the regions shown in panel A.

### Introgression analysis between H. cydno and H. melpomene

Previous studies suggest that introgression between *H. cydno* and its sister species *H. melpomene* has contributed to the diversification of mimetic wing patterns in this clade (Pardo-Diaz *et al*., 2012; Arias *et al*., 2012; Enciso-Romero *et al*., 2017; Thawornwattana *et al*., 2023). After identifying the candidate locus for the white submarginal band in *H. cydno*, we tested whether this region had introgressed into the *H. melpomene* subspecies exhibiting this trait (*H. m. cythera*), as previously suggested by Gilbert (2003).

Introgression was quantified with the fdM statistic (Martin *et al.,* 2015) across chromosome 15 and within the candidate region using Dsuite (Malinsky *et al*., 2021). fdM dynamically assigns the donor population, allowing for bidirectional gene flow while reducing variance and false positives in low-diversity regions (Martin *et al*., 2015). We defined four populations: (P1) *H. melpomene* subspecies lacking the white submarginal band; (P2) *H. m. cythera* with this band; (P3) *H. cydno* subspecies with the band; and (P4) *H. pardalinus* as outgroup (O). As a control, we repeated the analysis replacing P3 with *H. c. weymeri*, which lacks the trait and was expected to show fdM values near zero (Supplementary Fig. 2).

Sequences for *H. m. cythera, H. m. melpomene,* and *H. pardalinus* were downloaded from the European Nucleotide Archive (ENA), aligned to the same reference genome, and processed and filtered as described above (Supplementary Table 2). Dsuite’s *Dinvestigate* was run with sliding windows of 50 SNPs and a step size of 25 SNPs. Statistical support was assessed via permutation tests (*regioneR*; Gel *et al*., 2016), sampling 50,000 genomic regions of matching window size and comparing weighted mean fdM values in the candidate region against genome-wide expectations.

As additional evidence for introgression, we applied a topology-weighting approach with Twisst (Martin & Van Belleghem, 2017) to the same population configuration described above, testing clustering by phenotype (presence/absence of the white submarginal band), by phylogenetic relatedness, and a random topology. Species topology was defined as: (((*H. m. flavomartina + H. m. melpomene, H. m. cythera*)*, H. c. cydnides*)*, H. pardalinus*). ML trees were inferred with RAxML v8.2.9 (Stamatakis *et al*., 2008) in sliding windows of 50 SNPs (*raxml_sliding_windows.py*), retaining windows with ≥50% of individuals genotyped. Twisst was run with “--method complete” and default parameters.

## RESULTS

### Whole-genome resequencing data

For the 79 newly sequenced individuals, the average mapping rate to the reference genome was 95.83% (Supplementary Table 1). The proportion of missing data per individual ranged from 2% to 10%, with a few individuals showing 20–30% missing data; however, these individuals were retained because their coverage exceeded 7× (Supplementary Table 1). Mean depth of coverage ranged from 7× to 42× (Supplementary Table 1). The sequences generated for this paper have been submitted to the European Nucleotide Archive under the accession XXX-XXX.

### H. cydno subspecies tree and mtDNA network

The ASTRAL *H. cydno* subspecies tree, built from 248 ML gene trees, recovered the species as monophyletic with strong support and revealed two well-supported major clades corresponding to an east– west geographic split across the Colombian Central Cordillera (Fig. 1B), each further subdivided into subclades consistent with five geographic regions (Fig. 1A).

Within the eastern clade, each Venezuelan subspecies was monophyletic, and together they formed a monophyletic subclade (Figs. 1A, 1B). Magdalena Valley subspecies were paraphyletic (green), splitting into two subclades: one, sister to the Venezuelan samples, comprising *H. c. wanningeri*, *H. c. cydno*, and one individual of *H. c. hermogenes* (Figs. 1A, 1B); the second comprising *H. c. hermogenes* and *H. c. lisethae* together with several *H. c. cydno* individuals (Figs. 1A, 1B). Only *H. c. wanningeri* was recovered as monophyletic in this valley.

The western clade split into two internal clades. The first (orange) groups the Central American subspecies *H. c. galanthus* (monophyletic) and *H. c. chioneus* (paraphyletic), except for one individual from the Pacific (*H. c. zelinde*; Figs. 1A, 1B). The second clade comprises two subdivisions: the Cauca Valley subspecies *H. c. weymeri* and *H. c. cydnides* (purple), both reciprocally monophyletic with strong support; most *H. c. zelinde* individuals (light blue) together with *H. c. alithea* and *H. c. subsp. nov.* from Gorgona island, also both recovered as reciprocally monophyletic (Figs. 1A, 1B).

The mtDNA network reflected the same east–west split and partial geographic structure observed in the phylogenetic tree but showed multiple reticulations and no distinct clusters per subspecies (Fig. 1C). All *H. c. subsp. nov.* clustered together, as did nearly all *H. c. alithea* individuals (except for one sample, which grouped with the Gorgona cluster).

The eastern group formed one cluster of all Venezuelan individuals (pink), while Magdalena Valley samples were split across two clusters (green), both of which included Cauca Valley individuals (western group), and one was closely associated with Central American samples (orange; Figs. 1A, 1C). The western region showed three clusters: the least reticulated corresponded to Gorgona Island (dark blue) together with *H. c. alithea* (Figs. 1A, 1C); the other two included *H. c. weymeri* from the Cauca Valley, one grouping with *H. c. alithea* and the other with *H. c. zelinde* (Figs. 1A, 1C).

### Genetic variation and population structure

Consistent with the phylogenetic tree, clustering analyses recovered only 5 of the 14 described subspecies as distinct genetic units: *H. c. cydnides, H. c. weymeri, H. c. alithea, H. c. subsp. nov*. and *H. c. galanthus*, with an overall pattern consistent with geography. The first two PCA axes explained 29% of the variation and separated three geography-based clusters (Cauca Valley, Pacific, Gorgona Island), with a fourth cluster grouping Magdalena Valley, Venezuelan and Central American individuals (Fig. 2A). Because Gorgona individuals showed marked differentiation, we re-ran the PCA excluding this population to better resolve continental structure (Fig. 2B); this recovered the same groups.

**Figure 2.**
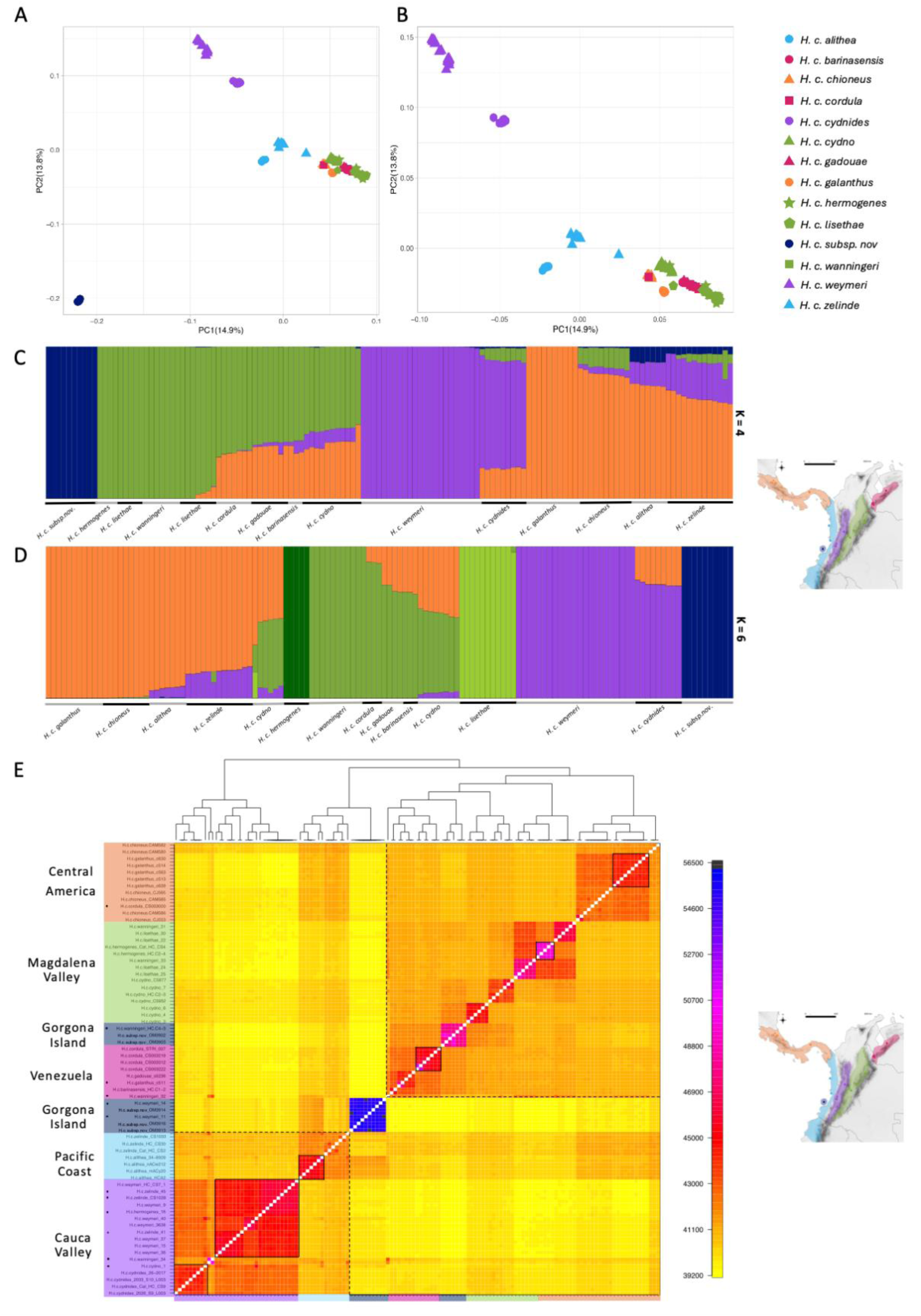
Genetic structure of *H. cydno* subspecies. **A)** PCA including all subspecies. **B)** PCA excluding Gorgona. The first two components explain 29% of the observed variation. **C)** ADMIXTURE plot for the best-fitting model (K = 4). **D)**fastSTRUCTURE plot for the best-fitting model (K = 6). Colors indicate geographic regions as shown in the map. **E)** FineSTRUCTURE matrix. The color scale indicates the level of shared genetic co-ancestry, from high (yellow) to low (blue), and the dendrogram shows genetic distances between groups. Black dots on the left indicate individuals assigned to genetic clusters outside their geographic region. Solid black boxes delimit individuals from the same subspecies, while dashed boxes indicate shared ancestry across geographic regions.

ADMIXTURE identified four genetic clusters as the best-fitting model (K = 4; AIC = 0.54785; Fig. 2C): Gorgona Island (dark blue), Magdalena Valley + Venezuela (green), Cauca Valley (purple), and Central America + Pacific (orange; Fig. 2C). Several subspecies showed marked heterogeneity in ancestry proportions; for example, some Magdalena Valley subspecies (e.g. *H. c. cydno*) shared ancestry with *H. c. weymeri* and the Central America–Pacific cluster, while *H. c. zelinde* drew ancestry from all four clusters. FastSTRUCTURE results were largely consistent with ADMIXTURE, though the best-supported model identified six clusters (maximum marginal likelihood; Fig. 2D): the same four recovered by ADMIXTURE, with the Magdalena Valley/Venezuela cluster further split into *H. c. lisethae* and *H. c. hermogenes* (Fig. 2D).

The FineSTRUCTURE coancestry matrix recovered two major clusters separating eastern and western regions (orange squares, Fig. 2E), with finer-scale structure within each: Central America clustered with the eastern populations of Venezuela and the Magdalena Valley. *H. c. subsp. nov.* from Gorgona formed a group outside the main clusters (blue region, Fig. 2E), though some individuals grouped with the eastern cluster (unoutlined pink square, Fig. 2E). Several highly structured, subspecies-exclusive clusters were also recovered (red regions, Fig. 2E): *H. c. cydnides, H. c. weymeri, H. c. alithea, H. c. cordula, H. c. hermogenes,* and *H. c. galanthus*. Some individuals were assigned to genetic clusters inconsistent with their geographic origin (black dots, Fig. 2E), reflecting extensive shared ancestry across regions; for example, one *H. c. hermogenes* individual clustered with Cauca Valley subspecies, and one *H. c. cordula* individual with the Central American individuals.

Pairwise genetic differentiation statistics among geographic regions and subspecies are shown in Supplementary Figs. 3A and 3B. Among regions, F_ST_ ranged from 0 to 0.4, with Gorgona Island the most structured (left, Supplementary Fig. 3A). Da showed a similar pattern, peaking at 0.06 between Gorgona and the Magdalena Valley (right, Supplementary Fig. 3A), while Dxy remained stable near 0.06 across regions (center, Supplementary Fig. 3A). Excluding Gorgona (given its outsized divergence) reduced maximum F_ST_ to 0.1, with the Cauca Valley now the most differentiated region (left, Supplementary Fig. 3B). Da showed a slightly different pattern, with Venezuela most structured in all pairwise comparisons (Da = 0.006; right, Supplementary Fig. 3B), while Dxy again remained stable (center, Supplementary Fig. 3B). At the subspecies level, excluding *H. c. subsp. nov.* from Gorgona Island (Supplementary Fig. 3C), the highest F_ST_ and Da values involved *H. c. alithea, H. c. wanningeri, H. c. weymeri,* and *H. c. cydnides*, while Dxy remained uniformly high (Supplementary Fig. 3D). Nucleotide diversity (π) was similar across subspecies with no clear geographic pattern, except for *H. c. subsp. nov.*, whose diversity was near zero (Supplementary Fig. 3E).

Genome-wide nucleotide diversity (π) and *D*xy showed a positive linear relationship (left, Supplementary Fig. 4), whereas *D*xy and *F*_ST_ did not: most windows had near-zero *F*_ST_ regardless of *D*xy (right, Supplementary Fig. 4). A subset of high-*D*xy, high-*F*_ST_ outlier windows mapped to chromosome 15, with additional outliers on chromosomes 20 and 21 (Z) in the parapatric comparisons *H. c. galanthus* vs. *H. c. chioneus*, *H. c. zelinde* vs. *H. c. alithea* and *H. c. cydno* vs. *H. c. hermogenes*, and in the allopatric comparison *H. c. wanningeri* and *H. c. alithea*.

### Origin of the radiation and demographic patterns

X- ORIGIN placed the most likely origin and expansion point of the radiation in northwestern Colombia, at the boundary between Central America (Panama) and South America (Colombia), in the Darién region (Fig. 3A).

**Figure 3.**
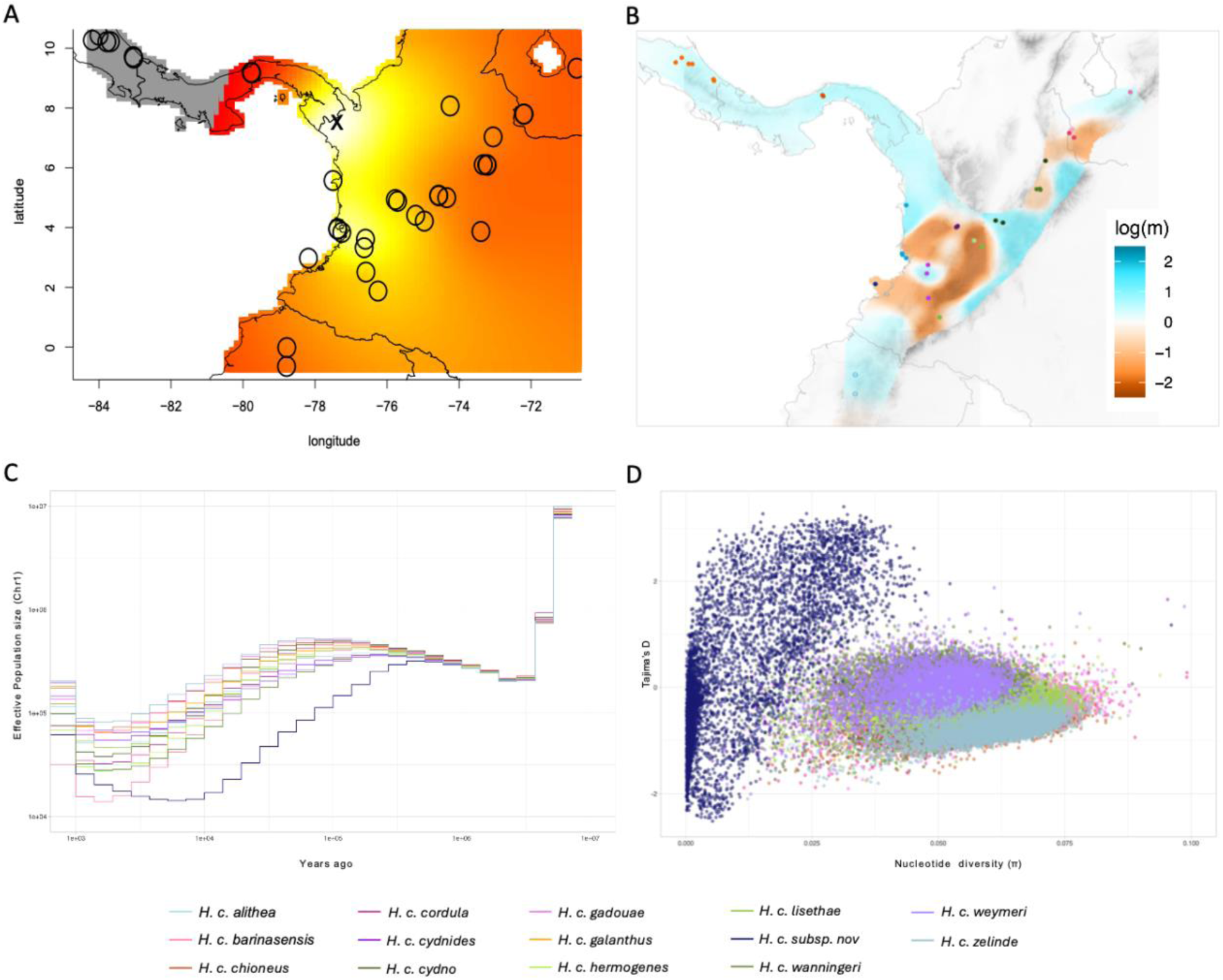
Geographic origin and demographic patterns of *H. cydno* subspecies. **A)** X-ORIGIN probability map of the geographic expansion. Darker tones indicate lower probabilities of origin, whereas lighter tones indicate higher probabilities. Circles represent the geographic coordinates of sampled individuals, and the “X” marks the location with the highest probability of origin. **B)** Migration surfaces inferred using EEMS. Dark blue regions indicate higher effective migration rates, whereas dark orange regions indicate lower genetic connectivity. **C)** Changes in effective population size through time inferred with Relate for chromosome 1. **D)** Tajima’s D values for each subspecies plotted against nucleotide diversity in 50kb windows.

EEMS revealed several areas with reduced migration. The lowest migration rates occurred in the Magdalena Valley, acting as an east–west barrier to gene flow, followed by the western slopes of the Western Cordillera, including parts of the Cauca Valley (dark orange, Fig. 3B). An additional migration barrier to gene flow was detected between the mainland and Gorgona Island, and in the southwestern of the Mérida Cordillera (Venezuela), extending to the western slope of the Eastern Cordillera (light orange, Fig. 3B). In contrast, high connectivity was observed between Central America and the northern Colombian Pacific, along northern Ecuador, across the eastern slope of the Andes, and in the northeastern Mérida Cordillera (blue, Fig. 3B).

Relate showed similar demographic patterns, characterized by a pronounced decline between 100,000 and 10,000 years ago and a subsequent gradual recovery toward the present. This pattern is illustrated for chromosome 1 (Fig. 3C) and was consistent across the remaining chromosomes (Supplementary Fig. 5). Notably, *H. c. subsp. nov.* exhibited an earlier and more pronounced decline in effective population size than continental subspecies, suggesting a population contraction potentially associated with a bottleneck. Tajima’s D plotted against nucleotide diversity was dispersed across subspecies. Most continental subspecies clustered near zero or in the negative range (0 to −1.5), whereas *H. c. subsp. nov.* showed more dispersed, predominantly positive values (−2 to 2), alongside near-zero nucleotide diversity, consistent with a recent population contraction followed by expansion, possibly linked to a founder effect (Fig. 3D).

### Genetic basis of the white submarginal band

The F_ST_ scan between *H. c. cydnides and H. c. weymeri,* which differ in all wing color bands and bars (Supplementary Fig. 1B), recovered strong signals that differ from the genomic background (F_ST_ > 0.4) at three previously documented color-pattern genes: *aristaless*, *WntA*, and *ivory* (Supplementary Fig. 6). We then focused on identifying the causal locus/loci for the unmapped white submarginal band.

Genome-wide F_ST_ scans grouping subspecies by phenotype identified elevated peaks on chromosomes 6, 20, and 21, but the strongest signal (F_ST_ > 0.2) was on chromosome 15 (Supplementary Fig. 7A), where a ∼500 Kb region ∼1.5 Mb from the chromosome start reached F_ST_ values near 0.3 (Supplementary Fig. 7B).

This genomic region was consistently recovered as differentiated (F_ST_ 0.5–0.7) across all four geographic comparisons (red shading, left, Fig. 4A; Supplementary Fig. 8) spanning positions *Hmel215003o*: 1,500,000–1,600,000 and containing the genes *Cortex* and *Parn*, the l*ncRNA ivory*, and two microRNAs, *miRNA193* and *miRNA2788* (red shading, right, Fig. 4A; Supplementary Fig. 8). In the Cauca Valley comparison, two additional peaks appeared upstream of the *ivory lncRNA* promoter (Supplementary Fig. 8B), coinciding with the yellow hindwing bar CREs *Yb(d)* and *Yb(v)* (Enciso-Romero *et al*., 2017). This was expected, as *H. c. weymeri* carries the yellow bar and *H. c. cydnides* does not.

**Figure 4.**
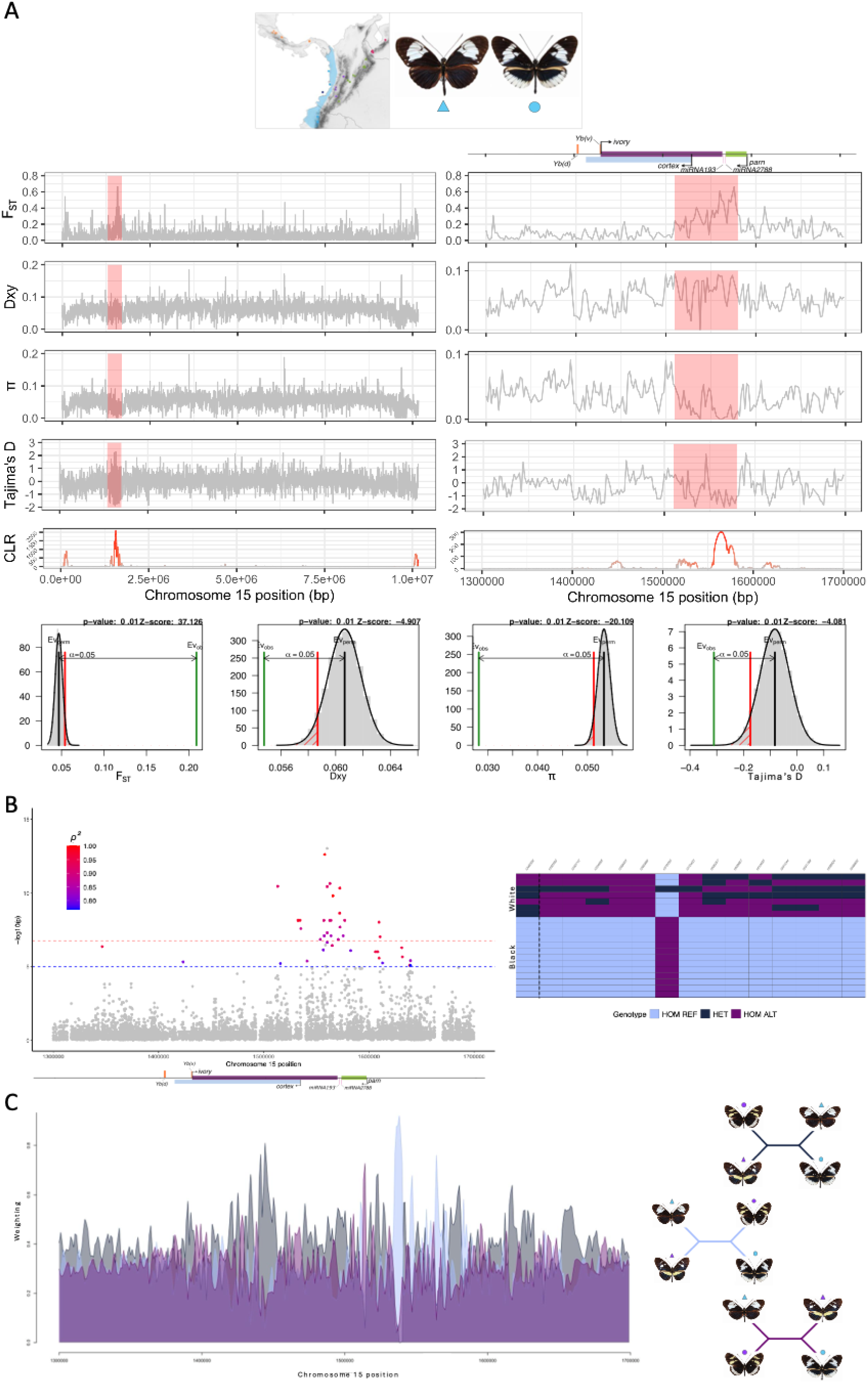
Genetic basis of the white submarginal band. **A)** Population differentiation, nucleotide diversity and signals of selection across chromosome 15 (left) and the candidate region (Hmel215003o: 1,500,000–1,600,000; right), showing relevant genes, CREs, miRNAs. The upper panel indicates the geographic region and phenotypes compared. The lower panel shows the randomization test results for each statistic across the genome, with the 5% significance threshold (red line) indicating the cutoff for the primary alternative hypothesis. In the case of selection, the color gradient indicates the estimated selection intensity (*α*), ranging from weak (grey) to strong (red) selection. **B)** GWAS of the *ivory:mir-193* locus and genotypes of the most strongly associated SNPs. The left panel shows Manhattan plots for the candidate region, with two significance thresholds: p-value = 1 × 10⁻⁵ (blue dashed line) and the Bonferroni-corrected threshold (red dashed line). SNPs exceeding both thresholds are colored according to the Spearman correlation between genotype and phenotype. The right panel shows reference homozygous, heterozygous, and alternate homozygous genotypes for the top-associated SNPs. Columns represent SNPs and rows represent individuals grouped by white submarginal band presence or absence. **C)** Twisst analysis of the Cauca Valley and Pacific regions. The left panel shows the topology weights across the candidate region. The right panel shows the three hypotheses evaluated: clustering by geographic origin (dark blue), phenotype (light blue), and the control hypothesis (purple).

Across all geographic comparisons, this genomic interval showed elevated Dxy and reduced π in subspecies with the white submarginal band (red shading, Fig. 4A; Supplementary Fig. 8), and a general trend toward negative Tajima’s D (reaching values near −2) most pronounced in the Pacific and Magdalena Valley + Venezuela comparisons (Fig. 4A; Supplementary Fig. 8C). All three statistics differed significantly from genome-wide null distributions based on permutations (Fig. 4A; Supplementary Fig. 8).

Consistent with the F_ST_ signal, GWAS peaks clustered within the terminal ivory lncRNA intron and the final exon, surrounding miRNA193, where SNPs showed the strongest genotype–phenotype association (Fig. 4B; Supplementary Fig. 9). The Cauca Valley comparison also showed peaks within the yellow hindwing bar CREs, coinciding with the F_ST_ signal there (Supplementary Fig. 9B). Fully associated SNPs clustered in the genomic interval *Hmel215003o*: 1,500,000–1,600,000 in all comparisons, showing haplotype blocks between phenotypic groups (Fig. 4B; Supplementary Fig. 9).

Twisst revealed phylogenetic discordance across the candidate region (Fig. 4C), and the geography-concordant topology predominated across most of the interval (dark blue, Fig. 4C). However, an alternative phenotype-concordant topology increased in frequency between positions 1,500,000–1,600,000 (light blue, Fig. 4C), consistent across all geographic comparisons (Supplementary Fig. 10) and supporting an association between this region and the white submarginal band.

SweepFinder2 provided strong, consistent support for positive selection on this candidate region across all geographic comparisons and both SFS null-model configurations (Fig. 4A; Supplementary Fig. 8), with CLR values ranging from 300 to 700. The highest CLR peaks coincided with the highest selection intensity (α) estimates. In the Cauca Valley comparison, additional selection signatures were detected within the yellow bar CREs, matching the F_ST_ and GWAS patterns (Supplementary Fig. 8B).

Given these concordant signals of genetic differentiation, association, and positive selection flanking *ivory*, we tested whether linkage disequilibrium (LD) maintains these loci together. LD between the candidate region associated with the white submarginal band and the regulatory elements associated with the yellow hindwing bar showed well-defined blocks of high LD (r² near 1, Supplementary Fig. 11). The first high-LD block was within *Yb(d)* and *Yb(v)* themselves, and the second spanned the interval between these elements and *miRNA193* (139,315 bp apart). This pattern reflects both physical linkage and the repulsion-phase association between the haplotypes underlying the two color traits in the hindwing.

### Signals of introgression between H. cydno and H. melpomene

Introgression analyses between *H. cydno* and *H. melpomene* (Supplementary Fig. 12) revealed elevated fdM values (∼0.5) on chromosome 15, indicating a strong signal of introgression between *H. c. cydnides* (white-banded) and *H. m. cythera* (Supplementary Fig. 12A). This pattern was also recovered using *H. c. alithea* (white-banded; Supplementary Fig. 13A). Replacing the banded subspecies with *H. c. weymeri* (band-absent) gave fdM values near zero across the same region (Supplementary Fig. 12B), showing no evidence of introgression.

Restricting the analysis to the candidate region (*Hmel215003o*: 1,500,000–1,600,000) showed a clear fdM peak when *H. c. cydnides* (Fig. 5A, purple) or *H. c. alithea* (Supplementary Fig. 13B) were included, versus a marked decrease for band-absent subspecies. The fdM maximum (red dashed line, Fig. 5A) fell within the terminal *ivory lncRNA* intron, upstream of *miRNA193*, and was significantly higher than expected under a null distribution generated by genome-wide permutations (Fig. 5B).

**Figure 5.**
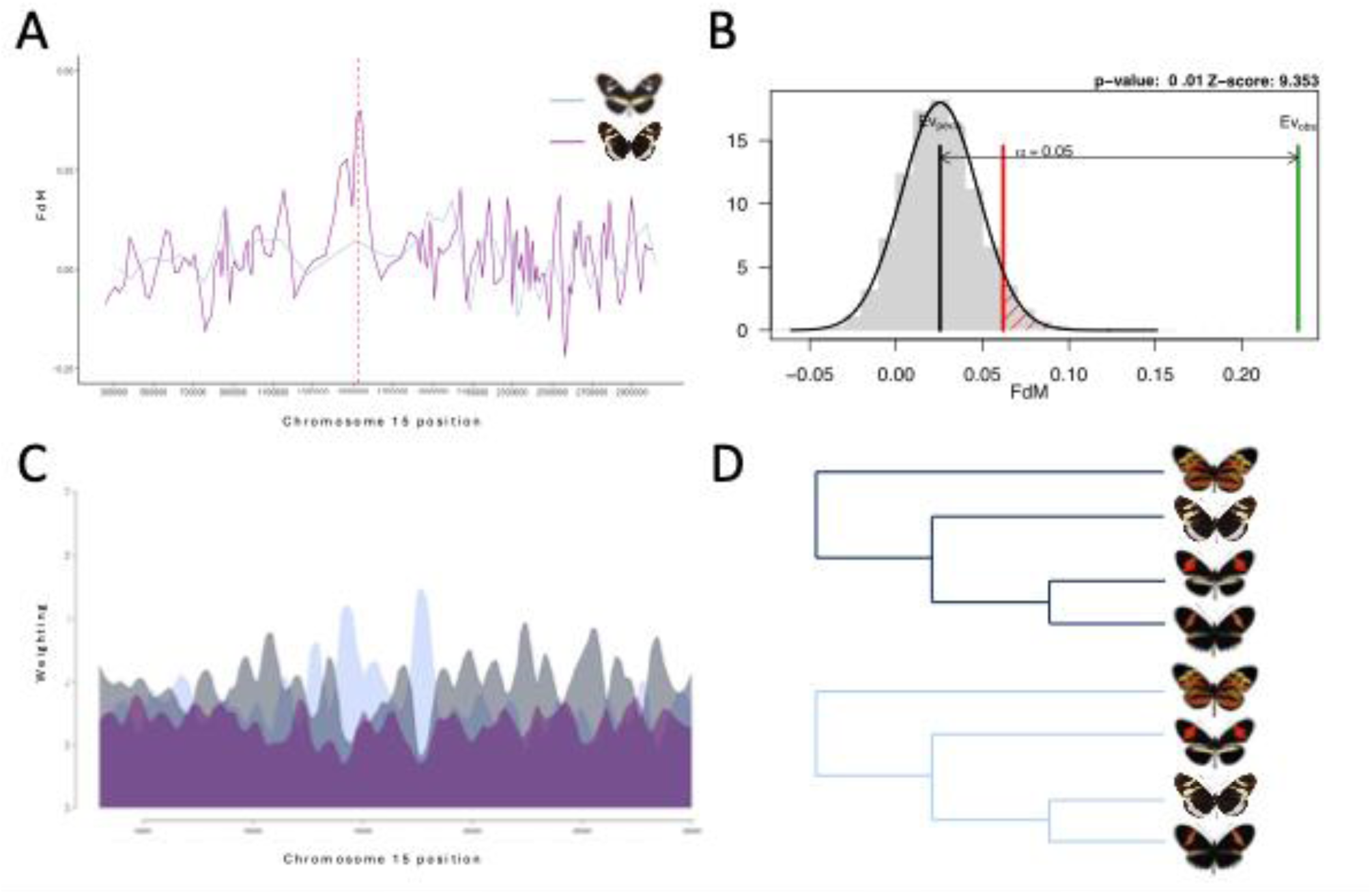
Introgression signals between *H. cydno* and *H. melpomene* at the *ivory:mir-193* locus. **A)** FdM statistics across the candidate region. The purple line represents the results for *H. c. cydnides*, which exhibits the white submarginal band, while the blue line represents the results for *H. c. weymeri,* which lacks this trait. See Supplementary Fig. 2 for the subspecies evaluated. **B)** Randomization test for FdM, with the 5% significance threshold (red line) indicating the cutoff for the primary alternative hypothesis. **C)** Twisst analysis results showing topology weights across the candidate region, colored by topology (left). The first topology represents the phylogenetic relationships (dark blue), with *H. m. cythera* grouped with *H. melpomene* (upper right panel). The second topology is associated with introgression of the white submarginal band phenotype (light blue), with *H. m. cythera* grouped with *H. cydno* (lower right panel). Purple indicates a random topology among these species (not shown).

As a second line of evidence, Twisst detected topological discordance across the region: the topology concordant with species/subspecies predominated overall, but at positions matching the fdM peak, an alternative topology grouping individuals by band increased in frequency (Fig. 5C and 5D; Supplementary Fig. 13C), further supporting introgression within the region spanning *ivory* and *miRNA193*.

## DISCUSSION

This study characterizes the phylogeography, genetic structure, and demographic history underlying the diversification of *Heliconius cydno*. This radiation comprises 14 morphologically and geographically defined subspecies distributed from Central America to northern South America, of which only nine represent distinct phylogenetic lineages: *H. c. barinasensis, H. c. cordula, H. c. gadouae, H. c. wanningeri, H. c. alithea, H. c. galanthus, H. c. subsp. nov., H. c. cydnides,* and *H. c. weymeri*. All nine formed monophyletic clades, and the latter five were consistently the most genetically structured groups across clustering analyses.

Four of these subspecies (*H. c. barinasensis, H. c. cordula, H. c. alithea*, and *H. c. wanningeri*) were also recovered by a previous AFLP-based study (Arias *et al*. 2014), but our whole-genome data resolved geographic structure with substantially greater phylogenetic resolution, likely because of the power of genome-wide data (Rannala & Yang, 2008; Davey *et al*., 2011; Andrews *et al*., 2016). Several subspecies were nonetheless paraphyletic in both studies, as expected under recent divergence, incomplete lineage sorting, or ongoing gene flow (Maddison, 1997; Degnan & Rosenberg, 2009; Toews & Brelsford, 2012). The genomic identification of only nine distinct lineages within *H. cydno* is relevant given the increasing importance of the subspecies category in lepidopteran evolution and conservation (Braby *et al*., 2012). We also found mito-nuclear discordance consistent with Arias *et al*. (2014), who reported paraphyletic and geographically intermixed subspecies based on the COI barcode. However, our complete mtDNA dataset recovered the east–west division described above (see Results), which the COI-based phylogeny lacked (Arias *et al*., 2014). Such nuclear and mitochondrial discordance is common across animal taxa (Toews & Brelsford, 2012; Hinojosa *et al*., 2019; Palacios *et al*., 2023; Salgado-Roa *et al*., 2024), including *Heliconius* and Ithomiini butterflies (Kozak *et al*., 2015; Van der Heijden *et al*., 2025).

Despite this discordance, the predominant geographic structure of *H. cydno* was consistent across datasets, likely reflecting the barrier effect of the Central Cordillera and the Magdalena Valley on gene flow. In fact, the Magdalena Valley is a documented barrier to connectivity in several taxa, though not previously in Lepidoptera, likely due to its steep precipitation gradient, highly heterogeneous forest types that range from dry to humid, and the Magdalena River itself (Gutiérrez-Pinto *et al*., 2012; Link *et al*., 2015; Muñoz-Ortiz *et al*., 2015; Sandoval-H *et al*., 2017; Chesser *et al*., 2020). The Central Cordillera is similarly an important barrier to gene flow in vertebrates and insects (Milá *et al*., 2009; Salgado-Roa *et al*., 2024), though its effect is not absolute, as species-specific dispersal can allow genetic exchange between populations on opposite sides (Smith *et al*., 2014). In this case, however, our results suggest limited east–west exchange in *H. cydno*. The high connectivity we observed within the Magdalena Valley itself (F_ST_ = 0.005–0.008, Supplementary Fig. 3D), where subspecies boundaries were poorly defined, contrasts with the local differentiation reported by Arias *et al*. (2014), suggesting no strong geographic, environmental, or ecological barriers to gene flow within the valley.

The Cauca Valley subspecies (*H. c. cydnides* in the north and *H. c. weymeri* in the south) show local genetic structure, with previous studies reporting significant mtDNA and nuclear differentiation and incipient asymmetric assortative mating (Arias *et al*., 2012). Our clustering and EEMS analyses likewise reveal a clear but partially permeable barrier to gene flow between the valley’s northern and southern portions, consistent with the previously characterized unimodal hybrid zone (∼15 km cline width) containing multi-generation hybrids and neutral-marker evidence of ongoing gene flow (Linares, 1997; Merchán *et al*., 2005; Arias *et al*., 2012). Our genome-wide F_ST_ estimate (0.05; Supplementary Fig. 3) was consistent with the estimate reported by Arias *et al*. (2012) based on dominant markers (AFLPs), but ∼4× lower than the microsatellite-based estimate (F_ST_ = 0.18) reported in the same study, likely reflecting differences in marker types and sampling. Furthermore, microsatellite-based estimates may be overestimated due to the presence of null alleles (Carlsson, 2008), while genome-wide estimates may dilute the contribution of highly differentiated loci associated with wing-pattern variation. Arias *et al*. (2012) attributed this genetic differentiation to a combination of ecological isolation, assortative mating, and forest-cover discontinuities rather than color pattern alone. Our results instead point to a stronger contribution from color-associated loci unlike Arias *et al*. (2014), whose AFLPs and microsatellites were not necessarily linked to color pattern genes. This is supported by elevated F_ST_ (∼0.5; Supplementary Fig. 6) at color pattern loci and by the fact that these subspecies belong to distinct mimicry rings (Brown, 1979; Linares, 1996, 1997). Consistently, Dxy and π showed a positive relationship (r = 0.95; Supplementary Fig. 4) across all pairwise comparisons, with greater dispersion among geographically distant subspecies and outlier loci carrying color pattern genes (chromosome 15), pheromone production genes (chromosome 20), and chromosome 21 (Z). The elevated differentiation on chromosome 21 may reflect its smaller effective population size as a sex chromosome.

Unlike the inter-Andean valleys, the northern range (Venezuela, Central America, and the Pacific) showed higher genetic connectivity and weaker structuring, suggesting greater gene flow across these regions via partially permeable mountain passes (Cadena *et al*., 2016), with local differentiation likely driven by geographic distance and mimicry-associated selection (Kapan, 2001; Chamberlain *et al*., 2009). *H. c. subsp. nov.* showed the strongest structure in the *H. cydno* radiation, consistent with a severe founder event following the colonization of Gorgona Island, which emerged ∼10–5 Ma from the Remolino–Grande–Gorgona structural high (41–23 Ma) and retained intermittent mainland connectivity thereafter (Pardo-Trujillo *et al*., 2020), likely allowing ancestral colonization during periods of lower sea levels before isolation by the current marine barrier.

Demographic history and dispersal patterns point to a *H. cydno* origin in the Darién region along the Colombia–Panama border, a biodiversity hotspot whose environmental complexity may have favored diversification (Myers *et al*., 2000; Kirby *et al*., 2008). This refines the broader northwestern South American/Central American origin proposed by *Arias et al.* (2014) and matches the inferred origin of *H. erato* in the same region (Quek *et al*., 2010; Hill *et al*., 2013; Guan *et al*. in prep.). In fact, the proposed origin of the radiation could explain the conflicting patterns of phylogenetic relationships and genetic clustering observed among Central American individuals. Their geographic proximity to this origin and historical connectivity may have preserved shared ancestral polymorphisms, although recent gene flow or a combination of both processes cannot be ruled out. All subspecies showed a marked reduction in effective population size between ∼100 and 10 Kya, potentially associated with the Last Glacial Period and the resulting contraction of tropical forests (Clark *et al*., 2009; Carnaval *et al*., 2009). However, this shared demographic signal could also reflect alternative processes such as mimetic advergence under shifting-balance dynamics (Mallet, 2010).

The origin of *H. cydn*o has been estimated at approximately 700 Kya (Guan *et al*., in prep.). By then, the major geological events shaping its current distribution had already occurred, including the final uplift of the Andes (∼5 Ma; Boschman, 2021; Hoorn *et al*., 2010), closure of the Isthmus of Panama (∼3 Ma; Coates & Obando, 1996; Kirby *et al*., 2008), and formation of the Magdalena (∼16–2.5 Ma; Sarmiento *et al*., 2015; Zapata *et al*., 2023) and Cauca (∼11.7–2.6 Ma; Padilla, 1981; López *et al*., 2005) valleys. This diversification history appears to have been driven primarily by dispersal rather than Andean vicariance, followed by local adaptation to regional mimicry patterns and assortative mating (Merrill *et al*., 2015; Cadena *et al*., 2016; Rodríguez-Muñoz *et al*., 2022; Salgado-Roa *et al*., 2024). Given the predominant role of dispersal, our results suggest multiple dispersal routes across the current range of *H. cydno*: northward into Central America via the Darién region and the Panamanian Isthmus (Kirby *et al*., 2008); southward through the Magdalena Valley and along the Pacific coast to Ecuador via the Chocó (Pérez-Gutiérrez, 2007); and toward Venezuela via the Táchira Depression (Haffer, 1969; Domínguez-Ossa, 1979; Cadena *et al*., 2016). Previously, Arias *et al*. (2014) proposed, from barcoding data alone, that northern and southern populations both colonized the Cauca Valley. Our mitochondrial genomes recover the same signal, but our nuclear data do not, suggesting local divergence followed by bidirectional gene flow between populations carrying distinct mitochondrial haplotypes.

For the genomic basis of the white submarginal hindwing band of *H. cydno*, we identified an association peak at positions 1,500,000 and 1,600,000 on chromosome 15, spanning the final intron and last exon of the *ivory* lncRNA, as well as *miRNA193,* which therefore represents the strongest candidate region for this phenotype and provides further evidence for the broader role of *ivory:mir-193* in lepidopteran wing color formation (Livraghi *et al*., 2021, 2024; Tian *et al*., 2024; Fandiño *et al*., 2024; Orteu *et al*., 2024; Ben-Chehida *et al*., 2026; Livraghi *et al*., 2026; Hanly *et al.,* 2026). *ivory* regulates melanic patterns via modulation of melanin-synthesis genes and controls white/yellow shifts in the absence of black pigmentation (Livraghi *et al*., 2024; Tian *et al*., 2024). Its first intron harbors cis-regulatory elements associated with seasonal color variation (Fandiño *et al*., 2024) and with yellow forewing band expression across Lepidoptera, contributing to phenotypic convergence (Ben-Chehida *et al*., 2026), while upstream and intragenic CREs are linked to white wing pattern variation (Orteu *et al*., 2024). Downstream of *ivory*, *miRNA193* is a major effector of melanic regulation (Tian *et al*., 2024), repressing melanin-synthesis genes and possibly regulating genes in the ommochrome pathway (Futahashi *et al*., 2021; Tian *et al*., 2024). Segregating cis-regulatory modules around *ivory* promote transcription of a lnc-pri-miRNA precursor that generates *miRNA193*, which itself acts as a co-transcriptional terminator (Livraghi *et al*., 2026; Hanly *et al.,* 2026), making *ivory* transcript and *miRNA193* a single regulatory unit. We therefore propose that *miRNA193* underlies variation in the white submarginal hindwing band across *H. cydno*, with reduced or altered *miRNA193* activity potentially reducing melanization and altering scale differentiation to produce the white band. The strong selective sweep signature at *miRNA193* between submarginal-banded and non-banded forms further suggests recent positive selection at this locus, potentially associated with mimetic advergence of *H. cydno* toward the *Heliconius sara*/*sapho* mimicry rings. Although we do not have functional validation, the evidence obtained in this study strongly supports the involvement of this locus, which has been repeatedly implicated in melanization across Lepidoptera. Nevertheless, the contribution of other genes to modify the phenotype should also be considered, such as WntA, which increases melanization in the distal wing veins along the hindwing margins (Concha *et al*., 2019).

Classical Mendelian studies identified linkage between *Yb* and *Sb*, with their alleles in repulsion (Linares, 1997; Merchán *et al*., 2005). *Yb* corresponds to the *Yb(d)/Yb(v)* regulatory modules upstream of *ivory* (Enciso-Romero *et al*., 2017; Livraghi *et al*., 2024), while our results identify *miRNA193* as the genetic basis of *Sb*. We found strong LD between *Yb(d)/Yb(v)* and *miRNA193* (r² > 0.8) despite their ∼139 Kb distance, with signatures of recent positive selection also reported at the *Yb* regulatory elements (Moest *et al.,* 2020). This flexible genetic architecture likely contributes to the maintenance of alternative mimetic patterns in the Cauca Valley, where recombinants carrying both traits occur at very low frequency within the hybrid zone between *H. c. cydnides* and *H. c. weymeri*. Furthermore, these color-pattern differences mediate asymmetric mating preferences between subspecies, with *H. c. cydnides* males showing a stronger incipient preference for their own color pattern than for that of *H. c. weymeri* (Arias *et al*., 2012).

Interspecific introgression is a recurrent source of adaptive variation in *Heliconius* (Edelman *et al*., 2019; Thawornwattana *et al*., 2023; Rossi *et al*., 2024), particularly at wing patterning loci such as *optix*, *ivory*, and the *Yb(d)/Yb(v)* regulatory elements (Pardo-Díaz *et al*., 2012; Dasmahapatra *et al*., 2012; Wallbank *et al*., 2016; Moest *et al*., 2020; Enciso-Romero *et al*., 2017). Our fdM and Twisst analyses reveal a further case of adaptive introgression between *H. cydno* and *H. melpomene* involving *ivory* and *miRNA193*. The most likely direction of transfer of the white submarginal band was from *H. cydno* to *H. m. cythera*, the only *H. melpomene* subspecies exhibiting this widespread *H. cydno* trait. Although previously proposed (Gilbert, 2003), our study provides the first quantitative evidence supporting this transfer, which may have facilitated a novel mimicry ring with *H. erato cyrbia*, illustrating how color pattern introgression can drive phenotypic diversification in *Heliconius*.

In conclusion, this study provides a comprehensive characterization of the phylogeography, genetic structure, gene flow, demographic history, and the genetic basis of wing color pattern variation in *H. cydno.* Genetic structure reflects both demographic history and recent selection associated with distinct mimicry rings, driving differentiation even amid gene flow among subspecies. We identify *ivory:mir-193* as the strongest candidate underlying variation in the mimetic submarginal hindwing band in *H. cydno*, a trait known as *Sb* locus but not previously mapped in *Heliconius*. As with other wing-pattern loci, we also found evidence of adaptive introgression at this locus, contributing to the formation of novel mimicry rings between *H. erato* and *H. melpomene*. This adds to the growing evidence that this locus facilitates adaptive convergence and divergence in Lepidoptera, in this case contributing to the formation of novel mimicry rings and incipient assortative mating between subspecies. More broadly, our results suggest that variation at phenotypic hotspots such as *ivory:mir-193* can generate intraspecific phenotypic diversity that, together with demographic processes, geographic connectivity, and local selection, contributes to rapid diversification and facilitates divergence during the early stages of adaptive radiation.

## ACKNOWLEDGMENTS

This work was supported by the COLCIENCIAS grant FP44842-103 -2015, awarded to C.S. Samples were collected under Permit No. 007, issued on October 7, 2015, by Parques Nacionales Naturales de Colombia to Universidad del Rosario for sample collection on Gorgona Island. All samples were collected under the framework of Environmental License No. 530 of 2014, issued by the National Environmental Licensing Authority (ANLA). We thank the HPC Service at Universidad del Rosario (CALDAS) and Sigma2, the National Infrastructure for High-Performance Computing and Data Storage in Norway, accessed through the University of Oslo, for providing access to computational resources. We also thank Dr. Juan Enciso-Romero and Dr. George Pacheco for their bioinformatics assistance.

