## Supporting Information for "Genomics of the diversification of *Heliconius cydno* (Lepidoptera: Nymphalidae) in northern South America: geographic and phenotypic patterns"

**Supplementary Table 1.** ***H. cydno*** **samples information and genotyping statistics.**

| Genus | Species | Race | ID | Locality | Latitude | Longitude | Hindwing phenotype | Mean depth per ind. | Missing data per ind. | % mapping genome | Coverage | Study accession | Source |
|---|---|---|---|---|---|---|---|---|---|---|---|---|---|
| *Heliconius* | *cydno* | *alithea* | 34-8509 | Pichincha, Ecuador | -0.6478 | -78.7894 | White submarginal band | 7.24 | 0.64 | 84.07 | 17.26 | PRJNA226620 | Gallant *et al.*(2014) |
| *Heliconius* | *cydno* | *alithea* | 35-8517 | Pichincha, Ecuador | -0.6478 | -78.7894 | White submarginal band | 13.10 | 0.28 | 93.45 | 18.85 | PRJNA226620 | Gallant *et al.* (2014) |
| *Heliconius* | *cydno* | *alithea* | HCA2 | Pichincha, Ecuador | -0.6478 | -78.7894 | White submarginal band | 17.50 | 0.06 | 96.78 | 19.11 | PRJNA226620 | Gallant *et al.* (2014) |
| *Heliconius* | *cydno* | *alithea* | mACw 172 | Pichincha, Ecuador | -0.0167 | -78.7833 | White submarginal band | 10.67 | 0.15 | 96.40 | 11.43 | PRJNA226620 | Gallant *et al.* (2014) |
| *Heliconius* | *cydno* | *alithea* | mACy 20 | Pichincha, Ecuador | -0.0167 | -78.7833 | White submarginal band | 9.81 | 0.15 | 96.40 | 11.43 | PRJNA226620 | Gallant *et al.* (2014) |
| *Heliconius* | *cydno* | *alithea* | nACw 212 | Pichincha, Ecuador | -0.0167 | -78.7833 | White submarginal band | 9.31 | 0.20 | 96.37 | 13.42 | PRJNA226620 | Gallant *et al.* (2014) |
| *Heliconius* | *cydno* | *alithea* | nACy112 | Pichincha, Ecuador | -0.0167 | -78.7833 | White submarginal band | 11.39 | 0.13 | 96.67 | 43.80 | PRJNA226620 | Gallant *et al.* (2014) |
| *Heliconius* | *cydno* | *barinasensis* | c1-1 | Trujillo, Venezuela | 9.3306 | -70.7311 | White submarginal band | 42.26 | 0.02 | 95.12 | 50.06 | | This study |
| *Heliconius* | *cydno* | *barinasensis* | c1-2 | Trujillo, Venezuela | 9.3306 | -70.7311 | White submarginal band | 26.59 | 0.03 | 93.61 | 30.70 | | This study |
| *Heliconius* | *cydno* | *chioneus* | CAM000580 | Colón, Panama | 9.1714 | -79.7572 | White submarginal band | 19.28 | 0.05 | 93.70 | 25.29 | PRJEB11772 | Martin *et al.* (2019) |

| | | | | | | | | | | | | | |
|---|---|---|---|---|---|---|---|---|---|---|---|---|---|
| *Heliconius* | *cydno* | *chioneus* | CAM000582 | Colón, Panama | 9.1714 | -79.7572 | White submarginal band | 21.48 | 0.04 | 94.42 | 22.05 | PRJEB11772 | Martin *et al.* (2019) |
| *Heliconius* | *cydno* | *chioneus* | CAM000586 | Colón, Panama | 9.1714 | -79.7572 | White submarginal band | 19.82 | 0.05 | 93.36 | 21.78 | PRJEB11772 | Martin *et al.* (2019) |
| *Heliconius* | *cydno* | *chioneus* | CAM584 | Colón, Panama | 9.1714 | -79.7572 | White submarginal band | 21.02 | 0.10 | 96.62 | 11.61 | PRJEB11772 | Van Belleghem *et al.* (2018) |
| *Heliconius* | *cydno* | *chioneus* | CAM585 | Colón, Panama | 9.1714 | -79.7572 | White submarginal band | 19.70 | 0.07 | 98.23 | 4.28 | PRJEB11772 | Van Belleghem *et al.* (2018) |
| *Heliconius* | *cydno* | *chioneus* | cyd553 | Colón, Panama | 9.1942 | -79.7728 | White submarginal band | 31.09 | 0.04 | 95.14 | 35.53 | ERP002440 | Martin *et al.* (2013) |
| *Heliconius* | *cydno* | *chioneus* | cyd560 | Colón, Panama | 9.1942 | -79.7728 | White submarginal band | 30.59 | 0.04 | 96.20 | 35.07 | ERP002440 | Martin *et al.* (2013) |
| *Heliconius* | *cydno* | *chioneus* | cyd564 | Colón, Panama | 9.1942 | -79.7728 | White submarginal band | 33.86 | 0.04 | 96.86 | 39.08 | ERP002440 | Martin *et al.* (2013) |
| *Heliconius* | *cydno* | *chioneus* | cyd565 | Colón, Panama | 9.1942 | -79.7728 | White submarginal band | 40.86 | 0.03 | 94.12 | 46.09 | ERP002440 | Martin *et al.* (2013) |
| *Heliconius* | *cydno* | *cordula* | c1-3 | San Cristóbal, Venezuela | 7.9406 | -72.3406 | Black hindwing | 23.73 | 0.03 | 94.12 | 26.57 | PRJEB21091 | Enciso-Romero *et al.* (2017) |
| *Heliconius* | *cydno* | *cordula* | c1-4 | San Cristóbal, Venezuela | 7.9406 | -72.3406 | Black hindwing | 23.31 | 0.03 | 93.73 | 27.66 | PRJEB21091 | Enciso-Romero *et al.* (2017) |
| *Heliconius* | *cydno* | *cordula* | CS003000 | San Cristóbal, Venezuela | 7.7939 | -72.1989 | Black hindwing | 18.41 | 0.03 | 97.32 | 31.28 | | This study |
| *Heliconius* | *cydno* | *cordula* | CS003218 | San Cristóbal, Venezuela | 7.7939 | -72.1989 | Black hindwing | 23.19 | 0.03 | 97.27 | 30.30 | | This study |

| | | | | | | | | | | | | |
|---|---|---|---|---|---|---|---|---|---|---|---|---|
| *Heliconius* | *cydno* | *cordula* | CS003 219 | San Cristóbal, Venezuela | 7.7939 | -72.1989 | Black hindwing | 18.87 | 0.04 | 96.65 | 22.82 | | This study |
| *Heliconius* | *cydno* | *cordula* | CS003 222 | San Cristóbal, Venezuela | 7.7939 | -72.1989 | Black hindwing | 23.32 | 0.03 | 96.51 | 32.62 | | This study |
| *Heliconius* | *cydno* | *cordula* | CS003 312 | San Cristóbal, Venezuela | 7.7939 | -72.1989 | Black hindwing | 23.86 | 0.04 | 87.74 | 27.46 | | This study |
| *Heliconius* | *cydno* | *cordula* | STRI-007 | San Cristóbal, Venezuela | 7.7933 | -72.1989 | Black hindwing | 34.44 | 0.03 | 85.05 | 12.56 | PRJEB8011 | Wallbank *et al.* (2016) |
| *Heliconius* | *cydno* | *cydnides* | 1569 | Risaralda, Colombia | 4.9400 | -75.7200 | White submarginal band | 31.75 | 0.02 | 96.78 | 35.92 | | This study |
| *Heliconius* | *cydno* | *cydnides* | 1600 | Cauca valley, Colombia | 3.8665 | -76.3834 | White submarginal band | 14.59 | 0.10 | 93.81 | 20.01 | | This study |
| *Heliconius* | *cydno* | *cydnides* | 2025 | Risaralda, Colombia | 4.8775 | -75.7103 | White submarginal band | 14.41 | 0.09 | 94.13 | 17.83 | | This study |
| *Heliconius* | *cydno* | *cydnides* | 2026 | Risaralda, Colombia | 4.8775 | -75.7103 | White submarginal band | 18.03 | 0.08 | 93.71 | 22.35 | | This study |
| *Heliconius* | *cydno* | *cydnides* | 2033 | Risaralda, Colombia | 4.8775 | -75.7103 | White submarginal band | 19.92 | 0.08 | 93.94 | 24.77 | | This study |
| *Heliconius* | *cydno* | *cydnides* | 2034 | Risaralda, Colombia | 4.8775 | -75.7103 | White submarginal band | 12.11 | 0.11 | 93.54 | 15.07 | | This study |
| *Heliconius* | *cydno* | *cydnides* | 26-2017 | Risaralda, Colombia | 4.8775 | -75.7103 | White submarginal band | 15.12 | 0.07 | 94.95 | 18.68 | PRJEB8011 | Wallbank *et al.* (2016) |
| *Heliconius* | *cydno* | *cydnides* | 27-2018 | Risaralda, Colombia | 4.8775 | -75.7103 | White submarginal band | 6.55 | 0.66 | 92.64 | 12.47 | PRJEB8011 | Wallbank *et al.* (2016) |
| *Heliconius* | *cydno* | *cydnides* | HC.CS 9 | Risaralda, Colombia | 4.8775 | -75.7103 | White submarginal band | 28.94 | 0.02 | 95.47 | 31.48 | | This study |
| *Heliconius* | *cydno* | *cydno* | 2076 | Bucaramanga, Colombia | 7.0317 | -73.0537 | White submarginal band | 9.19 | 0.14 | 96.62 | 10.24 | | This study |

| | | | | | | | | | | | | | |
|---|---|---|---|---|---|---|---|---|---|---|---|---|---|
| *Heliconius* | *cydno* | *cydno* | 2080 | Bucaramanga, Colombia | 7.0317 | -73.0537 | White submarginal band | 8.56 | 0.18 | 96.65 | 9.75 | | This study |
| *Heliconius* | *cydno* | *cydno* | 2141 | Bucaramanga, Colombia | 7.0317 | -73.0537 | White submarginal band | 9.47 | 0.14 | 96.49 | 10.40 | | This study |
| *Heliconius* | *cydno* | *cydno* | 2142 | Bucaramanga, Colombia | 7.0317 | -73.0537 | White submarginal band | 10.73 | 0.12 | 96.53 | 12.08 | | This study |
| *Heliconius* | *cydno* | *cydno* | 2223 | Cundinamarca, Colombia | 5.0792 | -74.5658 | White submarginal band | 10.79 | 0.10 | 96.43 | 12.23 | | This study |
| *Heliconius* | *cydno* | *cydno* | 3076 | Cundinamarca, Colombia | 5.0619 | -74.5658 | White submarginal band | 7.35 | 0.23 | 96.58 | 8.32 | | This study |
| *Heliconius* | *cydno* | *cydno* | 3079 | Cundinamarca, Colombia | 5.0619 | -74.5658 | White submarginal band | 8.08 | 0.20 | 96.58 | 8.32 | | This study |
| *Heliconius* | *cydno* | *cydno* | c2-2 | Santander, Colombia | 7.0317 | -73.0537 | White submarginal band | 12.46 | 0.14 | 94.51 | 41.72 | PRJEB21091 | Enciso-Romero *et al.* (2017) |
| *Heliconius* | *cydno* | *cydno* | c2-3 | Santander, Colombia | 7.0317 | -73.0537 | White submarginal band | 36.31 | 0.03 | 93.09 | 34.21 | PRJEB21091 | Enciso-Romero *et al.* (2017) |
| Heliconius | *cydno* | *cydno* | C5877 | Cundinamarca, Colombia | 4.9992 | -74.3411 | White submarginal band | 15.86 | 0.08 | 97.65 | 22.20 | | This study |
| Heliconius | *cydno* | *cydno* | C5890 | Cundinamarca, Colombia | 4.9992 | -74.3411 | White submarginal band | 15.94 | 0.04 | 97.51 | 15.40 | | This study |
| Heliconius | *cydno* | *cydno* | C5912 | Cundinamarca, Colombia | 4.9992 | -74.3411 | White submarginal band | 15.39 | 0.11 | 95.32 | 32.01 | | This study |
| Heliconius | *cydno* | *cydno* | C5952 | Cundinamarca, Colombia | 4.9992 | -74.3411 | White submarginal band | 10.87 | 0.10 | 66.59 | 41.05 | | This study |

| | | | | | | | | | | | | | |
|---|---|---|---|---|---|---|---|---|---|---|---|---|---|
| *Heliconius* | *cydno* | *gadouae* | 5233 | Táchira, Venezuela | 8.0687 | -72.2480 | Black hindwing | 15.77 | 0.05 | 98.17 | 22.36 | | This study |
| *Heliconius* | *cydno* | *gadouae* | 5234 | Táchira, Venezuela | 8.0687 | -72.2480 | Black hindwing | 17.65 | 0.04 | 97.90 | 23.18 | | This study |
| *Heliconius* | *cydno* | *gadouae* | 5235 | Táchira, Venezuela | 8.0687 | -72.2480 | Black hindwing | 19.03 | 0.05 | 97.88 | 13.37 | | This study |
| *Heliconius* | *cydno* | *gadouae* | 5236 | Táchira, Venezuela | 8.0687 | -72.2480 | Black hindwing | 16.18 | 0.04 | 98.07 | 12.85 | | This study |
| *Heliconius* | *cydno* | *gadouae* | 5237 | Táchira, Venezuela | 8.0687 | -72.2480 | Black hindwing | 16.85 | 0.04 | 98.18 | 16.53 | | This study |
| *Heliconius* | *cydno* | *galanthus* | c511 | Cariblanco, Costa Rica | 10.2667 | -84.1833 | Black hindwing | 15.31 | 0.07 | 94.46 | 24.81 | SRA106228 | Kronforst *et al.* (2013) |
| *Heliconius* | *cydno* | *galanthus* | c512 | Limón, Costa Rica | 9.6667 | -83.0333 | Black hindwing | 15.40 | 0.07 | 93.84 | 30.87 | SRA106228 | Kronforst *et al.* (2013) |
| *Heliconius* | *cydno* | *galanthus* | c513 | Heredia, Costa Rica | 10.4333 | -84.0000 | Black hindwing | 15.66 | 0.07 | 94.42 | 13.28 | SRA106228 | Kronforst *et al.* (2013) |
| *Heliconius* | *cydno* | *galanthus* | c514 | Vesta, Costa Rica | 9.7167 | -83.0500 | Black hindwing | 15.29 | 0.08 | 94.48 | 33.02 | SRA106228 | Kronforst *et al.* (2013) |
| *Heliconius* | *cydno* | *galanthus* | c515 | Guacimo, Costa Rica | 10.2167 | -83.6833 | Black hindwing | 15.21 | 0.08 | 94.26 | 25.56 | SRA106228 | Kronforst *et al.* (2013) |
| *Heliconius* | *cydno* | *galanthus* | c563 | Guapiles, Costa Rica | 10.2167 | -83.7833 | Black hindwing | 14.61 | 0.08 | 94.61 | 28.58 | SRA106228 | Kronforst *et al.* (2013) |
| *Heliconius* | *cydno* | *galanthus* | c614 | Vesta, Costa Rica | 9.7167 | -83.0500 | Black hindwing | 15.35 | 0.07 | 94.70 | 27.18 | SRA106228 | Kronforst *et al.* (2013) |
| *Heliconius* | *cydno* | *galanthus* | c630 | Heredia, Costa Rica | 10.4333 | -84.0000 | Black hindwing | 14.46 | 0.08 | 94.21 | 15.71 | SRA106228 | Kronforst *et al.* (2013) |
| *Heliconius* | *cydno* | *galanthus* | c639 | Guacimo, Costa Rica | 10.2167 | -83.6833 | Black hindwing | 15.33 | 0.08 | 94.47 | 16.77 | SRA106228 | Kronforst *et al.* (2013) |

| | | | | | | | | | | | | | |
|---|---|---|---|---|---|---|---|---|---|---|---|---|---|
| *Heliconius* | *cydno* | *galanthus* | c640 | Vesta, Costa Rica | 9.7167 | -83.0500 | Black hindwing | 15.10 | 0.08 | 94.25 | 15.74 | SRA106228 | Kronforst *et al.* (2013) |
| *Heliconius* | *cydno* | *hermogenes* | 3112 | Tolima, Colombia | 4.4092 | -75.2061 | White submarginal band | 10.27 | 0.10 | 96.74 | 11.20 | | This study |
| *Heliconius* | *cydno* | *hermogenes* | 3116 | Tolima, Colombia | 4.4092 | -75.2061 | White submarginal band | 9.44 | 0.12 | 96.63 | 10.20 | | This study |
| *Heliconius* | *cydno* | *hermogenes* | c2-1 | Tolima, Colombia | 4.4092 | -75.2061 | White submarginal band | 29.59 | 0.03 | 94.51 | 14.62 | | This study |
| *Heliconius* | *cydno* | *hermogenes* | c2-4 | Tolima, Colombia | 4.4092 | -75.2061 | White submarginal band | 24.89 | 0.04 | 93.97 | 29.23 | | This study |
| *Heliconius* | *cydno* | *hermogenes* | HC.CS 3 | Tolima, Colombia | 4.4092 | -75.2061 | White submarginal band | 32.34 | 0.02 | 96.20 | 35.07 | | This study |
| *Heliconius* | *cydno* | *hermogenes* | HC.CS 4 | Tolima, Colombia | 4.4092 | -75.2061 | White submarginal band | 28.78 | 0.03 | 96.86 | 39.08 | | This study |
| *Heliconius* | *cydno* | *lisethae* | 2371 | Huila, Colombia | 1.8736 | -76.2524 | White submarginal band | 10.68 | 0.10 | 96.66 | 12.13 | | This study |
| *Heliconius* | *cydno* | *lisethae* | 2372 | Huila, Colombia | 1.8736 | -76.2524 | White submarginal band | 11.21 | 0.11 | 96.84 | 12.80 | | This study |
| *Heliconius* | *cydno* | *lisethae* | 2377 | Huila, Colombia | 1.8736 | -76.2524 | White submarginal band | 10.83 | 0.11 | 96.63 | 12.41 | | This study |
| *Heliconius* | *cydno* | *lisethae* | 2380 | Huila, Colombia | 1.8736 | -76.2524 | White submarginal band | 9.67 | 0.12 | 96.70 | 10.54 | | This study |
| *Heliconius* | *cydno* | *lisethae* | 2383 | Huila, Colombia | 1.8736 | -76.2524 | White submarginal band | 10.86 | 0.12 | 96.40 | 12.36 | | This study |
| *Heliconius* | *cydno* | *lisethae* | 3378 | Huila, Colombia | 1.8736 | -76.2524 | White submarginal band | 8.68 | 0.23 | 96.31 | 10.03 | | This study |

| | | | | | | | | | | | | | |
|---|---|---|---|---|---|---|---|---|---|---|---|---|---|
| *Heliconius* | *cydno* | *lisethae* | 3379 | Huila, Colombia | 1.8736 | -76.2524 | White submarginal band | 9.00 | 0.17 | 96.56 | 10.13 | | This study |
| *Heliconius* | *cydno* | *lisethae* | 3382 | Huila, Colombia | 1.8736 | -76.2524 | White submarginal band | 9.07 | 0.15 | 96.58 | 10.05 | | This study |
| *Heliconius* | *cydno* | *lisethae* | 3431 | Tolima, Colombia | 4.2147 | -74.9653 | White submarginal band | 9.76 | 0.12 | 96.70 | 10.70 | | This study |
| *Heliconius* | *cydno* | *lisethae* | c3-1 | Tolima, Colombia | 4.2147 | -74.9653 | White submarginal band | 21.13 | 0.06 | 94.59 | 16.66 | | This study |
| *Heliconius* | *cydno* | *lisethae* | c3-2 | Tolima, Colombia | 4.2147 | -74.9653 | White submarginal band | 24.93 | 0.05 | 94.07 | 16.83 | | This study |
| *Heliconius* | *cydno* | *wanningeri* | c4-2/2963 | Santander, Colombia | 6.1014 | -73.3286 | White submarginal band | 9.02 | 0.03 | 94.54 | 16.79 | | This study |
| *Heliconius* | *cydno* | *wanningeri* | c4-3/2963 | Santander, Colombia | 6.1014 | -73.3286 | White submarginal band | 10.26 | 0.05 | 94.42 | 16.68 | | This study |
| *Heliconius* | *cydno* | *wanningeri* | 1589 | Santander, Colombia | 6.0856 | -73.2186 | White submarginal band | 10.94 | 0.17 | 96.38 | 10.02 | | This study |
| *Heliconius* | *cydno* | *wanningeri* | 2062 | Santander, Colombia | 6.1022 | -73.2286 | White submarginal band | 9.89 | 0.12 | 96.54 | 11.28 | | This study |
| *Heliconius* | *cydno* | *wanningeri* | 2954 | Santander, Colombia | 6.1022 | -73.2286 | White submarginal band | 8.70 | 0.11 | 96.38 | 12.17 | | This study |
| *Heliconius* | *cydno* | *wanningeri* | 2955 | Santander, Colombia | 6.1022 | -73.2286 | White submarginal band | 28.63 | 0.13 | 96.60 | 10.93 | | This study |
| *Heliconius* | *cydno* | *wanningeri* | 2956 | Santander, Colombia | 6.1022 | -73.2286 | White submarginal band | 21.96 | 0.19 | 96.14 | 9.83 | | This study |
| *Heliconius* | *cydno* | *subsp. nov.* | 3899 | Gorgona island, Colombia | 2.9731 | -78.1847 | Black hindwing | 16.42 | 0.07 | 97.44 | 18.73 | | This study |

| | | | | | | | | | | | | | |
|---|---|---|---|---|---|---|---|---|---|---|---|---|---|
| *Heliconius* | *cydno* | *subsp. nov.* | 3900 | Gorgona island, Colombia | 2.9731 | -78.1847 | Black hindwing | 14.31 | 0.08 | 97.08 | 16.19 | | This study |
| *Heliconius* | *cydno* | *subsp. nov.* | 3901 | Gorgona island, Colombia | 2.9731 | -78.1847 | Black hindwing | 15.79 | 0.07 | 97.02 | 17.97 | | This study |
| *Heliconius* | *cydno* | *subsp. nov.* | 3902 | Gorgona island, Colombia | 2.9731 | -78.1847 | Black hindwing | 14.00 | 0.08 | 97.72 | 15.86 | | This study |
| *Heliconius* | *cydno* | *subsp. nov.* | 3903 | Gorgona island, Colombia | 2.9731 | -78.1847 | Black hindwing | 16.21 | 0.07 | 97.77 | 18.33 | | This study |
| *Heliconius* | *cydno* | *subsp. nov.* | 3912 | Gorgona island, Colombia | 2.9731 | -78.1847 | Black hindwing | 16.50 | 0.07 | 97.43 | 18.60 | | This study |
| *Heliconius* | *cydno* | *subsp. nov.* | 3913 | Gorgona island, Colombia | 2.9731 | -78.1847 | Black hindwing | 11.82 | 0.11 | 97.60 | 20.30 | | This study |
| *Heliconius* | *cydno* | *subsp. nov.* | 3914 | Gorgona island, Colombia | 2.9731 | -78.1847 | Black hindwing | 11.28 | 0.12 | 97.02 | 22.09 | | This study |
| *Heliconius* | *cydno* | *subsp. nov.* | 3915 | Gorgona island, Colombia | 2.9731 | -78.1847 | Black hindwing | 13.88 | 0.16 | 94.47 | 25.41 | | This study |
| *Heliconius* | *cydno* | *subsp. nov.* | 3916 | Gorgona island, Colombia | 2.9731 | -78.1847 | Black hindwing | 10.77 | 0.14 | 97.04 | 27.60 | | This study |
| *Heliconius* | *cydno* | *weymeri f. weymeri* | HC.CS 6 | Cauca valley, Colombia | 3.3242 | -76.6364 | Black hindwing | 52.60 | 0.02 | 94.12 | 46.09 | PRJEB21091 | Enciso-Romero *et al.* (2017) |
| *Heliconius* | *cydno* | *weymeri f. weymeri* | HC.CS 7 | Cauca valley, Colombia | 3.3242 | -76.6364 | Black hindwing | 41.81 | 0.02 | 97.32 | 31.28 | PRJEB21091 | Enciso-Romero *et al.* (2017) |
| *Heliconius* | *cydno* | *weymeri f. weymeri* | HC.CS 8 | Cauca valley, Colombia | 3.3242 | -76.6364 | Black hindwing | 30.87 | 0.02 | 97.27 | 30.30 | PRJEB21091 | Enciso-Romero *et al.* (2017) |
| *Heliconius* | *cydno* | *weymeri f. weymeri* | STRI00 61 | Cauca valley, Colombia | 3.3242 | -76.6364 | Black hindwing | 26.34 | 0.04 | 96.65 | 22.82 | PRJEB21091 | Enciso-Romero *et al.* (2017) |
| *Heliconius* | *cydno* | *weymeri f. weymeri* | STRI00 63 | Cauca valley, Colombia | 3.3242 | -76.6364 | Black hindwing | 16.78 | 0.06 | 96.51 | 32.62 | PRJEB21091 | Enciso-Romero *et al.* (2017) |

| | | | | | | | | | | | | | |
|---|---|---|---|---|---|---|---|---|---|---|---|---|---|
| *Heliconius* | *cydno* | *weymeri f. gustavi* | 1956 | Cauca valley, Colombia | 3.6100 | -76.5994 | Black hindwing | 7.58 | 0.23 | 96.52 | 8.39 | | This study |
| *Heliconius* | *cydno* | *weymeri f. gustavi* | 2507 | Cauca valley, Colombia | 3.6100 | -76.5994 | Black hindwing | 7.83 | 0.21 | 96.39 | 8.71 | | This study |
| *Heliconius* | *cydno* | *weymeri f. gustavi* | 2508 | Cauca valley, Colombia | 3.6100 | -76.5994 | Black hindwing | 7.08 | 0.30 | 96.44 | 8.10 | | This study |
| *Heliconius* | *cydno* | *weymeri f. gustavi* | 2517 | Cauca valley, Colombia | 3.6100 | -76.5994 | Black hindwing | 7.42 | 0.29 | 96.40 | 8.44 | | This study |
| *Heliconius* | *cydno* | *weymeri f. gustavi* | 2518 | Cauca valley, Colombia | 3.6100 | -76.5994 | Black hindwing | 8.76 | 0.17 | 96.50 | 9.73 | | This study |
| *Heliconius* | *cydno* | *weymeri f. gustavi* | 2519 | Cauca valley, Colombia | 3.6100 | -76.5994 | Black hindwing | 9.91 | 0.13 | 96.54 | 10.89 | | This study |
| *Heliconius* | *cydno* | *weymeri f. gustavi* | 2522 | Cauca valley, Colombia | 3.6100 | -76.5994 | Black hindwing | 10.35 | 0.11 | 96.52 | 11.39 | | This study |
| *Heliconius* | *cydno* | *weymeri f. gustavi* | 2538 | Cauca valley, Colombia | 3.3242 | -76.6364 | Black hindwing | 8.84 | 0.18 | 96.54 | 9.91 | | This study |
| *Heliconius* | *cydno* | *weymeri f. gustavi* | 2904 | Cauca valley, Colombia | 3.3242 | -76.6364 | Black hindwing | 9.58 | 0.14 | 96.17 | 10.64 | | This study |
| *Heliconius* | *cydno* | *weymeri f. gustavi* | 3630 | Cauca valley, Colombia | 3.3242 | -76.6364 | Black hindwing | 10.66 | 0.12 | 96.69 | 11.89 | | This study |
| *Heliconius* | *cydno* | *weymeri f. gustavi* | 3633 | Cauca valley, Colombia | 3.3242 | -76.6364 | Black hindwing | 9.27 | 0.14 | 96.29 | 10.35 | | This study |
| *Heliconius* | *cydno* | *weymeri f. gustavi* | 3634 | Cauca valley, Colombia | 3.3242 | -76.6364 | Black hindwing | 11.85 | 0.09 | 96.42 | 13.22 | | This study |
| *Heliconius* | *cydno* | *weymeri f. gustavi* | 29-2529 | Cauca valley, Colombia | 2.5081 | -76.5867 | Black hindwing | 6.62 | 0.43 | 92.10 | 10.70 | PRJEB8011 | Wallbank *et al.* (2016) |
| *Heliconius* | *cydno* | *weymeri f. gustavi* | M3635 | Cauca valley, Colombia | 3.3242 | -76.6364 | Black hindwing | 31.09 | 0.03 | 87.74 | 27.46 | PRJEB21091 | Enciso-Romero *et al.* (2017) |
| *Heliconius* | *cydno* | *weymeri f. gustavi* | M3638 | Cauca valley, Colombia | 3.3242 | -76.6364 | Black hindwing | 27.12 | 0.04 | 95.54 | 24.33 | PRJEB21091 | Enciso-Romero *et al.* (2017) |
| *Heliconius* | *cydno* | *weymeri f. gustavi* | M3640 | Cauca valley, Colombia | 3.3242 | -76.6364 | Black hindwing | 36.90 | 0.03 | 95.09 | 26.75 | PRJEB21091 | Enciso-Romero |

| | | | | | | | | | | | | | |
|---|---|---|---|---|---|---|---|---|---|---|---|---|---|
| | | | | | | | | | | | | | *et al.* (2017) |
| *Heliconius* | *cydno* | *weymeri f. gustavi* | M3637 | Cauca valley, Colombia | 3.3242 | -76.6364 | Black hindwing | 28.69 | 0.03 | 96.78 | 19.11 | PRJEB21091 | Enciso-Romero *et al.* (2017) |
| *Heliconius* | *cydno* | *weymeri f. gustavi* | M3639 | Cauca valley, Colombia | 3.3242 | -76.6364 | Black hindwing | 29.12 | 0.03 | 95.14 | 35.53 | PRJEB21091 | Enciso-Romero *et al.* (2017) |
| *Heliconius* | *cydno* | *zelinde* | 3490 | Cauca valley, Colombia | 3.8369 | -77.2573 | Black hindwing | 9.09 | 0.17 | 96.69 | 10.01 | | This study |
| *Heliconius* | *cydno* | *zelinde* | 3493 | Cauca valley, Colombia | 3.8369 | -77.2573 | Black hindwing | 10.74 | 0.10 | 96.80 | 11.63 | | This study |
| *Heliconius* | *cydno* | *zelinde* | 3530 | Chocó, Colombia | 5.5715 | -77.5017 | Black hindwing | 10.19 | 0.10 | 96.28 | 11.10 | | This study |
| *Heliconius* | *cydno* | *zelinde* | 32-2262 | Cauca valley, Colombia | 3.9394 | -77.3689 | Black hindwing | 6.65 | 0.35 | 92.25 | 9.32 | PRJEB11772 | Martin *et al.* (2019) |
| *Heliconius* | *cydno* | *zelinde* | CS001035 | Cauca valley, Colombia | 3.9394 | -77.3689 | Black hindwing | 27.28 | 0.02 | 96.75 | 34.30 | PRJEB11772 | Martin *et al.* (2019) |
| *Heliconius* | *cydno* | *zelinde* | CS1028 | Cauca valley, Colombia | 3.9394 | -77.3689 | Black hindwing | 30.52 | 0.02 | 95.16 | 35.41 | PRJEB11772 | Martin *et al.* (2019) |
| *Heliconius* | *cydno* | *zelinde* | CS1029 | Cauca valley, Colombia | 3.9583 | -77.3733 | Black hindwing | 21.11 | 0.05 | 97.92 | 28.18 | PRJEB11772 | Martin *et al.* (2019) |
| *Heliconius* | *cydno* | *zelinde* | CS1030 | Cauca valley, Colombia | 3.9583 | -77.3733 | Black hindwing | 22.95 | 0.06 | 97.76 | 30.74 | PRJEB11772 | Martin *et al.* (2019) |
| *Heliconius* | *cydno* | *zelinde* | CS1033 | Cauca valley, Colombia | 3.9394 | -77.3689 | Black hindwing | 24.29 | 0.05 | 98.12 | 32.47 | PRJEB11772 | Martin *et al.* (2019) |
| *Heliconius* | *cydno* | *zelinde* | CS273 | Cauca valley, Colombia | 3.9394 | -77.3689 | Black hindwing | 24.08 | 0.03 | 94.92 | 28.03 | PRJEB11772 | Martin *et al.* (2019) |
| *Heliconius* | *cydno* | *zelinde* | HC.CS1 | Cauca valley, Colombia | 3.9394 | -77.3689 | Black hindwing | 32.75 | 0.03 | 95.74 | 36.60 | PRJEB21091 | Enciso-Romero *et al.* (2017) |
| *Heliconius* | *cydno* | *zelinde* | HC.CS2 | Cauca valley, Colombia | 3.9394 | -77.3689 | Black hindwing | 31.72 | 0.02 | 95.37 | 33.98 | PRJEB21091 | Enciso-Romero *et al.* (2017) |

| | | | | | | | | | | | | | |
|---|---|---|---|---|---|---|---|---|---|---|---|---|---|
| *Heliconius* | *cydno* | *zelinde* | HC.CS 30 | Cauca valley, Colombia | 3.9394 | -77.3689 | Black hindwing | 39.18 | 0.02 | 95.89 | 42.74 | PRJEB21091 | Enciso-Romero *et al.* (2017) |

**Supplementary Table 2. *Heliconius* species used as outgroups and *H. melpomene* subspecies included in the Dsuite analysis, with their accession numbers.**

| Genus | Species | Race | ID | Study accession | Source |
|---|---|---|---|---|---|
| *Heliconius* | *melpomene* | *malleti* | HMMal.CS21 | PRJEB8011 | Wallbank *et al.* (2016) |
| *Heliconius* | *melpomene* | *malleti* | HMMal.CS22 | PRJEB8011 | Wallbank *et al.* (2016) |
| *Heliconius* | *melpomene* | *malleti* | HMMal.CS24 | PRJEB8011 | Wallbank *et al.* (2016) |
| *Heliconius* | *melpomene* | *malleti* | HMMalS.CS000482 | PRJEB8011 | Wallbank *et al.* (2016) |
| *Heliconius* | *melpomene* | *malleti* | HMMalS.CS004030 | PRJEB8011 | Wallbank *et al.* (2016) |
| *Heliconius* | *melpomene* | *malleti* | HMMalS.CS004034 | PRJEB8011 | Wallbank *et al.* (2016) |
| *Heliconius* | *melpomene* | *melpomene* | HMMcol.CS25 | PRJEB11772 | Martin *et al.* (2019) |
| *Heliconius* | *melpomene* | *melpomene* | HMMcol.CS26 | PRJEB11772 | Martin *et al.* (2019) |
| *Heliconius* | *melpomene* | *melpomene* | HMMcol.CS27 | PRJEB11772 | Martin *et al.* (2019) |
| *Heliconius* | *melpomene* | *melpomene* | HMMfg.CAM2035 | PRJEB11772 | Martin *et al.* (2019) |
| *Heliconius* | *melpomene* | *melpomene* | HMMfg.CAM8216 | PRJEB11772 | Martin *et al.* (2019) |
| *Heliconius* | *melpomene* | *melpomene* | HMMfg.CAM8218 | PRJEB11772 | Martin *et al.* (2019) |
| *Heliconius* | *melpomene* | *melpomene* | HMMpan.16-18097 | ERP002440 | Martin *et al.* (2013) |
| *Heliconius* | *melpomene* | *melpomene* | HMMpan.17-18038 | ERP002440 | Martin *et al.* (2013) |
| *Heliconius* | *melpomene* | *melpomene* | HMMpan.CAM014673 | PRJEB11772 | Martin *et al.* (2019) |
| *Heliconius* | *melpomene* | *flavomartina* | HMR.4169 | | This study |
| *Heliconius* | *melpomene* | *flavomartina* | HMR.4201 | | This study |
| *Heliconius* | *melpomene* | *flavomartina* | HMR.4228 | | This study |
| *Heliconius* | *melpomene* | *cythera* | HM.cythera.CAM040382 | PRJEB35570 | Montejo-Kovacevich *et al.* (2022) |
| *Heliconius* | *melpomene* | *cythera* | HM.cythera.CAM040383 | PRJEB35570 | Montejo-Kovacevich *et al.* (2022) |
| *Heliconius* | *melpomene* | *cythera* | HM.cythera.CAM040456 | PRJEB35570 | Montejo-Kovacevich *et al.* (2022) |
| *Heliconius* | *melpomene* | *cythera* | HM.cythera.CAM040457 | PRJEB35570 | Montejo-Kovacevich *et al.* (2022) |
| *Heliconius* | *melpomene* | *cythera* | HM.cythera.CAM040458 | PRJEB35570 | Montejo-Kovacevich *et al.* (2022) |
| *Heliconius* | *melpomene* | *cythera* | HM.cythera.CAM040459 | PRJEB35570 | Montejo-Kovacevich *et al.* (2022) |
| *Heliconius* | *melpomene* | *cythera* | HM.cythera.CAM040460 | PRJEB35570 | Montejo-Kovacevich *et al.* (2022) |
| *Heliconius* | *melpomene* | *cythera* | HM.cythera.CAM040475 | PRJEB35570 | Montejo-Kovacevich *et al.* (2022) |
| *Heliconius* | *melpomene* | *cythera* | HM.cythera.CAM040516 | PRJEB35570 | Montejo-Kovacevich *et al.* (2022) |
| *Heliconius* | *melpomene* | *cythera* | HM.cythera.CAM040517 | PRJEB35570 | Montejo-Kovacevich *et al.* (2022) |
| *Heliconius* | *melpomene* | *cythera* | HM.cythera.CAM040518 | PRJEB35570 | Montejo-Kovacevich *et al.* (2022) |
| *Heliconius* | *melpomene* | *cythera* | HM.cythera.CAM040519 | PRJEB35570 | Montejo-Kovacevich *et al.* (2022) |
| *Heliconius* | *timareta* | *florencia* | HTF.CS12 | PRJEB11772 | Martin *et al.* (2019) |
| *Heliconius* | *timareta* | *florencia* | HTF.CS13 | PRJEB11772 | Martin *et al.* (2019) |
| *Heliconius* | *timareta* | *florencia* | HTF.CS14 | PRJEB11772 | Martin *et al.* (2019) |
| *Heliconius* | *timareta* | *thelxinoe* | HTTh.Hel11-84 | ERP002440 | Martin *et al.* (2013) |
| *Heliconius* | *timareta* | *thelxinoe* | HTTh.Hel2-86 | ERP002440 | Martin *et al.* (2013) |
| *Heliconius* | *timareta* | *thelxinoe* | HTTh.Hel6-313 | ERP002440 | Martin *et al.* (2013) |
| *Heliconius* | *pardalinus* | *sergestus* | ERR1143615 | PRJEB8011 | Wallbank *et al.* (2016) |
| *Heliconius* | *pardalinus* | *sergestus* | ERR1143616 | PRJEB8011 | Wallbank *et al.* (2016) |
| *Heliconius* | *pardalinus* | *sergestus* | ERR1143617 | PRJEB8011 | Wallbank *et al.* (2016) |
| *Heliconius* | *pardalinus* | *sergestus* | ERR1143618 | PRJEB8011 | Wallbank *et al.* (2016) |
| *Heliconius* | *pardalinus* | *sergestus* | ERR1143619 | PRJEB8011 | Wallbank *et al.* (2016) |

| | | | | | |
|---|---|---|---|---|---|
| *Heliconius* | *pardalinus* | *sergestus* | ERR1143620 | PRJEB8011 | Wallbank *et al.* (2016) |
| *Heliconius* | *pardalinus* | - | SRR7162654 | PRJNA471310 | Jay *et al.* (2018) |
| *Heliconius* | *pardalinus* | - | SRR7162657 | PRJNA471310 | Jay *et al.* (2018) |
| *Heliconius* | *pardalinus* | - | SRR7162658 | PRJNA471310 | Jay *et al.* (2018) |
| *Heliconius* | *pardalinus* | - | SRR7162645 | PRJNA471310 | Jay *et al.* (2018) |
| *Heliconius* | *pardalinus* | - | SRR7162646 | PRJNA471310 | Jay *et al.* (2018) |
| *Heliconius* | *pardalinus* | - | SRR7162653 | PRJNA471310 | Jay *et al.* (2018) |

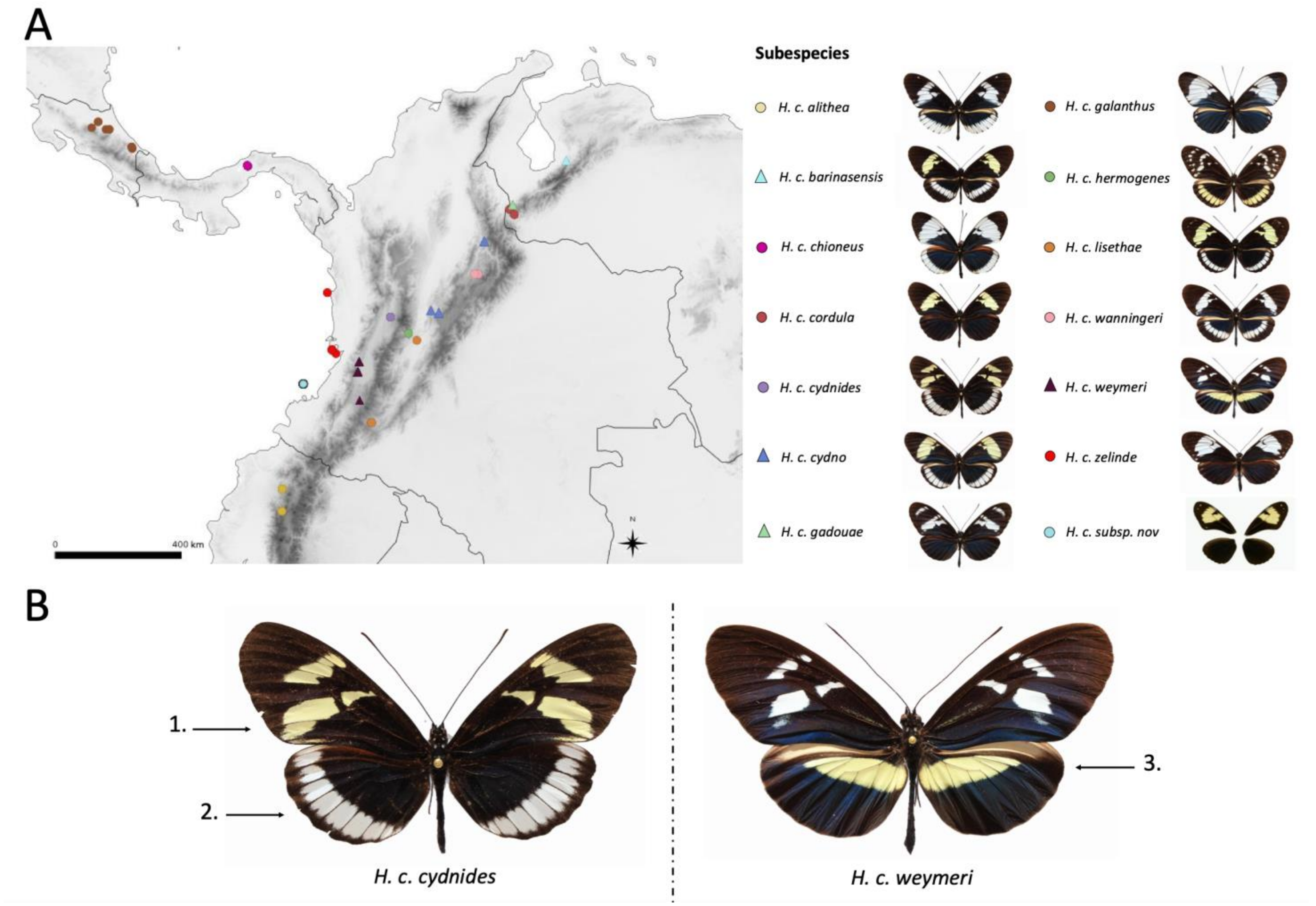


**Supplementary figure 1. Sampling distribution and color patterns of *H. cydno* subspecies. A)** Sampling map of the *H. cydno* subspecies included in this study. Symbols indicate sampling localities and corresponding phenotypes. Information for each sampling locality is provided in Supplementary Table 1. **B)** Color patterns of *H. c. cydnides* and *H. c. weymeri.* 1. White/yellow bands in the medial area of the forewing. 2. White/Black submarginal hindwing band. 3. Black/yellow hindwing bar.

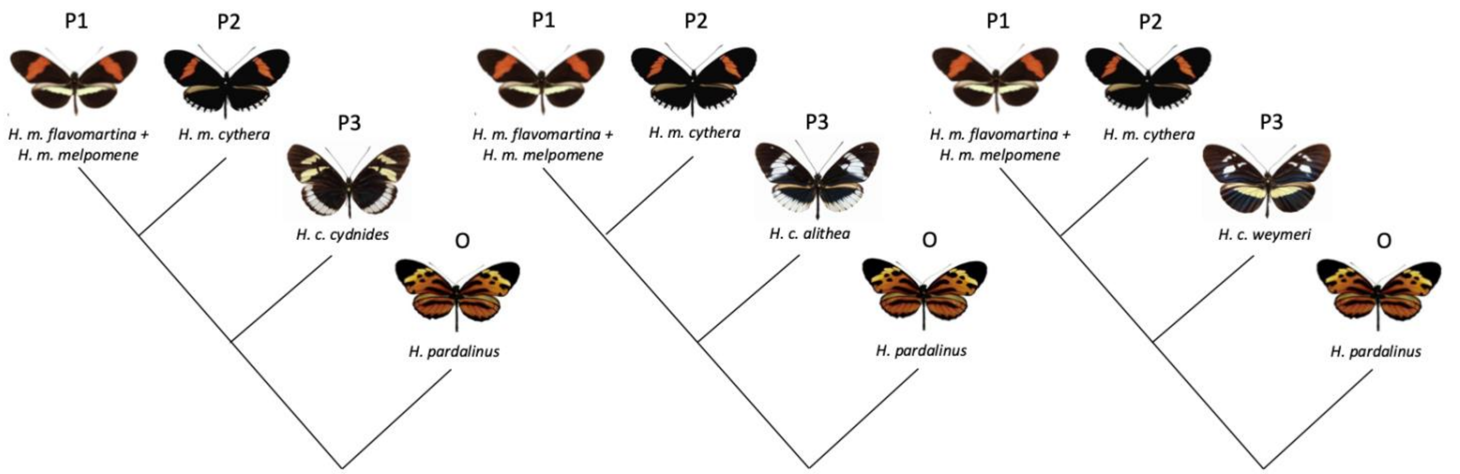


**Supplementary Figure 2. Population groupings used for introgression analyses quantified with the fdM statistic.** The left panel shows the first analysis, with *H. melpomene* subspecies lacking the white submarginal band (P1), *H. m. cythera* with the band (P2), *H. cydno cydnides* with the band (P3), and *H. pardalinus* as the outgroup (O). The central panel shows the second analysis, with *H. melpomene* (P1), *H. m. cythera* (P2), *H. cydno alithea* (P3), and *H. pardalinus* as the outgroup (O). The right panel shows the third analysis, with *H. melpomene* (P1), *H. m. cythera* (P2), *H. cydno weymeri* (P3), and *H. pardalinus* as the outgroup (O).

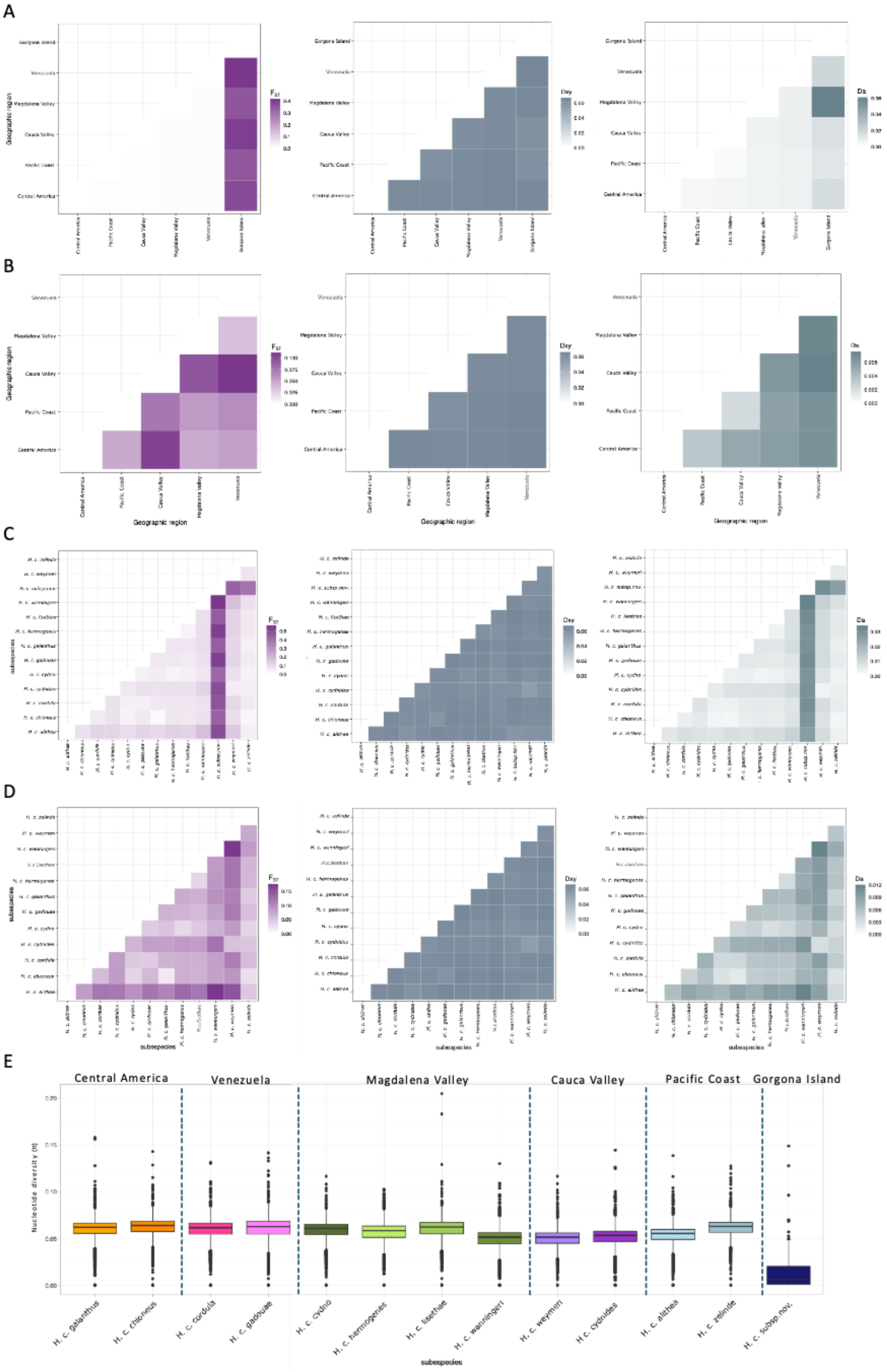


**Supplementary figure 3. Genetic differentiation and diversity statistics. A)** $F_{ST}$, Dxy, and Da between geographic regions. **B)** Same as A, excluding Gorgona. **C)** $F_{ST}$, Dxy, and Da between subspecies. **D)** Same as C, excluding Gorgona individuals. In all panels, purple heatmaps correspond to $F_{ST}$, blue heatmaps to Dxy, and green heatmaps to Da. **E)** Nucleotide diversity (π) boxplots for each subspecies. Boxes represent the first and third quartiles, and whiskers extend to the largest and smallest values within 1.5 × IQR.

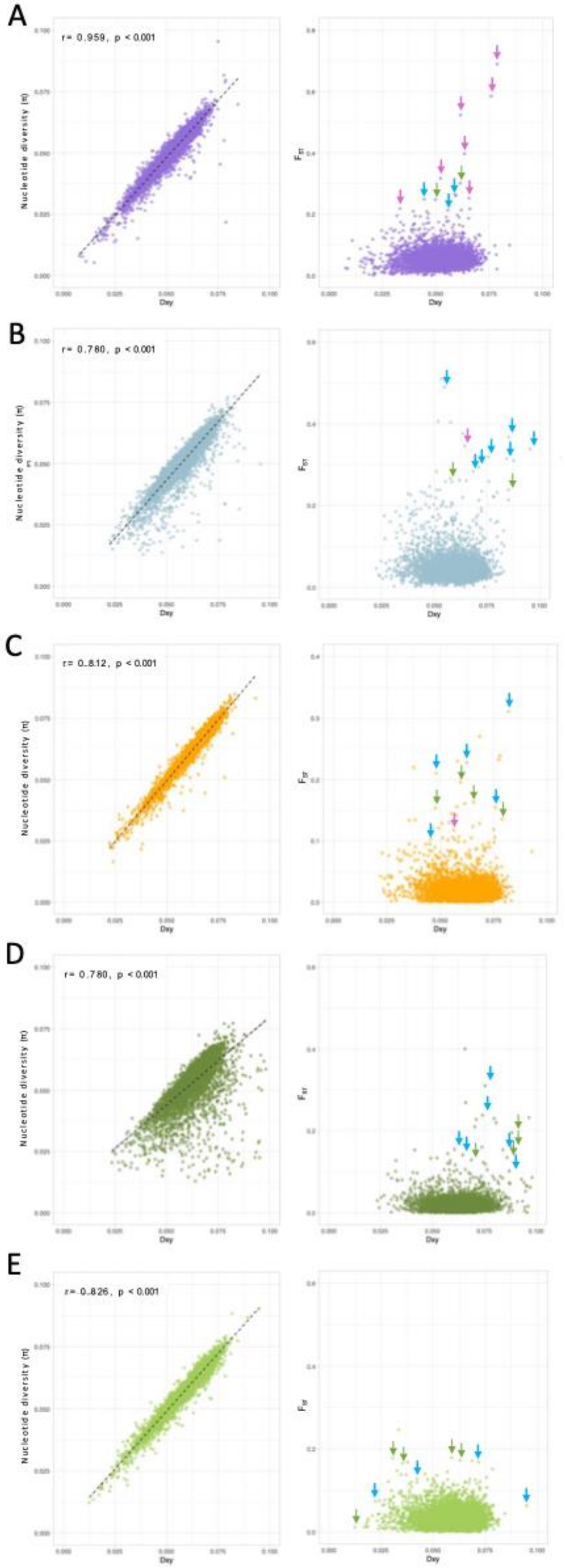


**Supplementary figure 4. Scatter plots of π versus Dxy (left panels) and $F_{ST}$ versus Dxy (right panels) calculated across the genome**. Pink arrows indicate outliers on chromosome 15, green arrows indicate outliers on chromosome 21, and blue arrows indicate outliers on chromosome 20. r represents the Pearson correlation coefficient. **A)** *H. c. cydnides* × *H. c. weymeri* **B)** *H. c. alithea* × *H. c. zelinde* **C)** *H. c. chioneus* × *H. c. galanthus* **D)** *H. c. alithea* × *H. c. wanningeri* **E)** *H. c. cydno* × *H. c. hermogenes*.

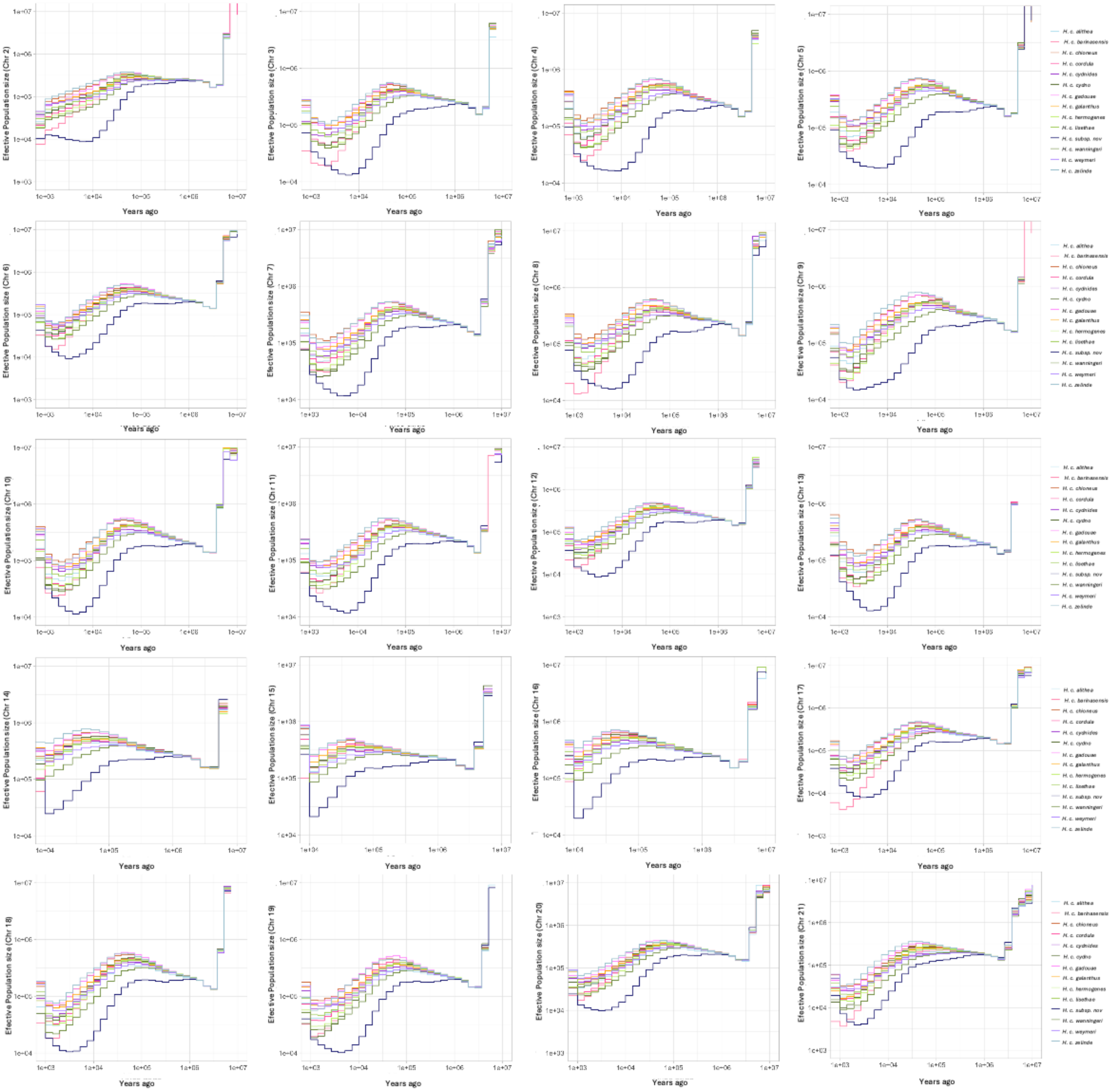


**Supplementary figure 5. Changes in effective population size through time inferred with Relate for each chromosome.** A consistent pattern of Ne fluctuations was observed across all subspecies and chromosomes.

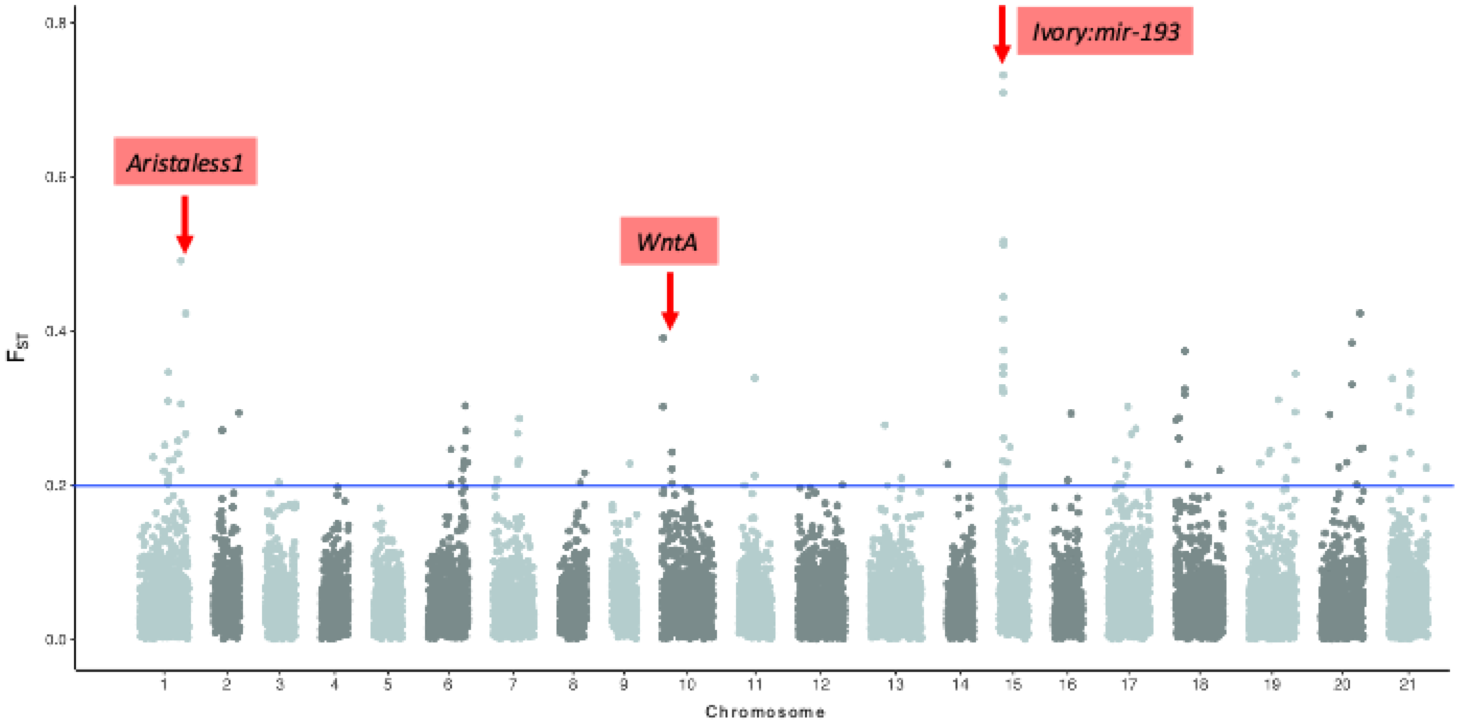


**Supplementary Figure 6. Genome-wide genetic differentiation between *H. c. cydnides* and *H. c. weymeri***. Red arrows indicate $F_{ST}$ peaks corresponding to loci previously associated with color pattern that differ between the two subspecies. The blue line indicates the significance threshold (p-value = 0.01).

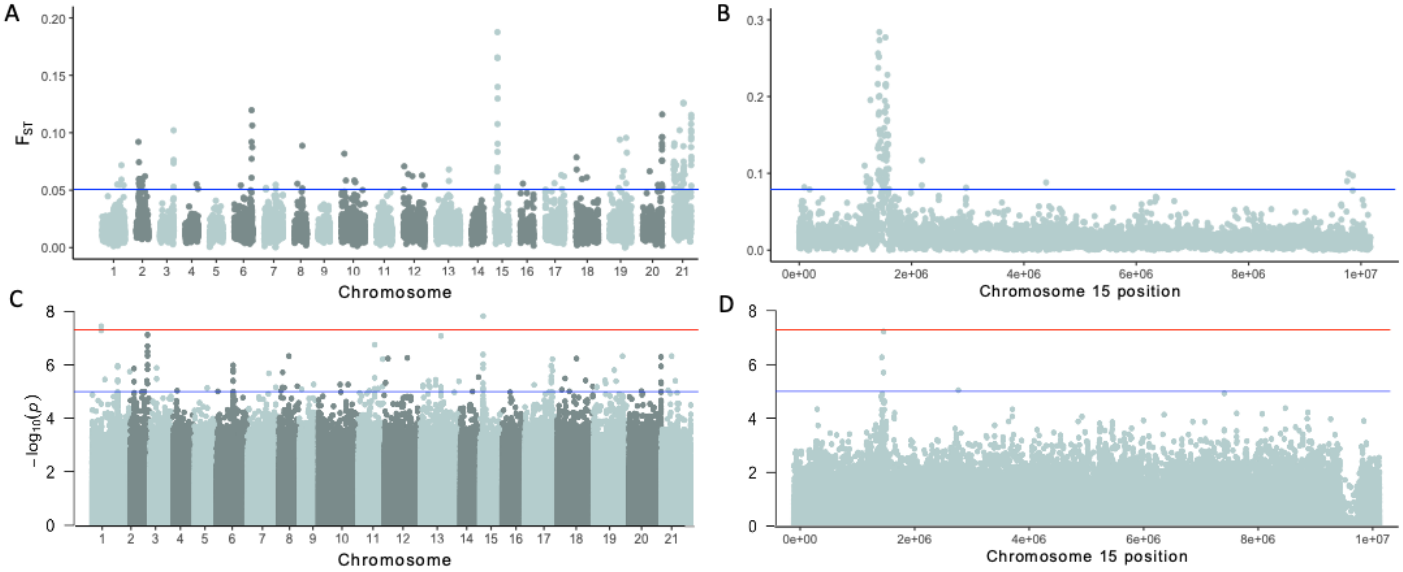


**Supplementary Figure 7. Genome-wide genetic differentiation and association analysis in *H. cydno***. The analysis includes all subspecies grouped according to their phenotype. **A)** Genome-wide $F_{ST}$ values calculated in non-overlapping 50 kb windows. **B)** $F_{ST}$ values across chromosome 15 calculated in non-overlapping 2 kb windows. The blue line indicates the significance threshold (p-value = 0.01). **C)** Genome-wide GWAS results. **D)** GWAS results for chromosome 15. Manhattan plots show two significance thresholds: p-value = $1 \times 10^{-5}$ (blue line) and the Bonferroni-corrected threshold (red line).

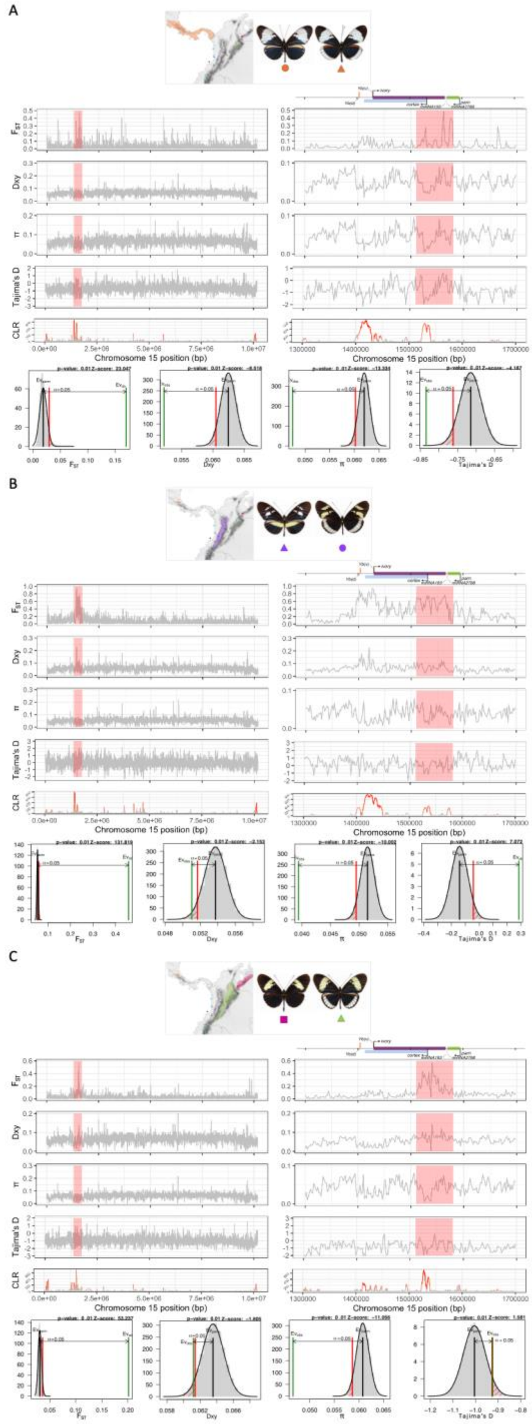


**Supplementary Figure 8. Genome scans across chromosome 15 and the candidate region by geographic region.** The left panel shows values across the entire chromosome 15 and the right panel shows values across the candidate region (Hmel215003o: 1,500,000–1,600,000), showing relevant genes, CREs, miRNAs. The upper panel indicates the geographic region and phenotypes compared. The lower panel shows the randomization test results for each statistic across the genome, with the 5% significance threshold (red line) indicating the cutoff for the primary alternative hypothesis. In the case of selection, the color gradient indicates the estimated selection intensity ($\alpha$), ranging from weak (grey) to strong (red) selection. **A)** Central America. **B)** Cauca Valley. **C)** Magdalena Valley and Venezuela.

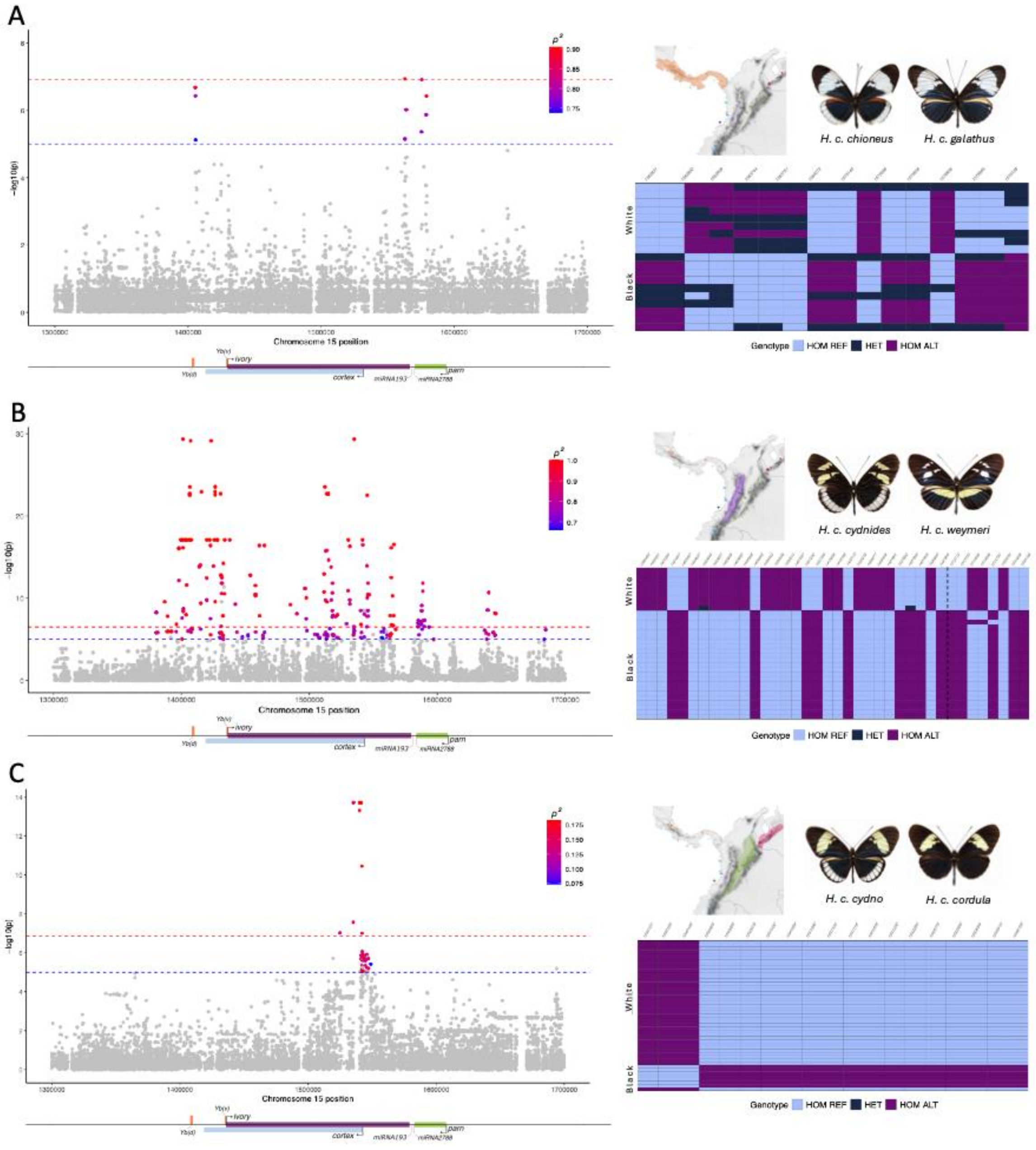


**Supplementary Figure 9**. **Genome-wide association study (GWAS) of the *ivory:mir-193* locus and genotypes of the most strongly associated SNPs by geographic region.** The left panel shows Manhattan plots for the candidate region (Hmel215003o: 1,500,000–1,600,000), alongside the positions of relevant genes, CREs, and miRNAs. The left panel shows Manhattan plots for the candidate region, with two significance thresholds: p-value = $1 \times 10^{-5}$ (blue dashed line) and the Bonferroni-corrected threshold (red dashed line). SNPs exceeding both thresholds are colored according to the Spearman correlation between genotype and phenotype. The right panel shows the geographic region, phenotypes compared, and genotypes (reference homozygous, heterozygous, and alternate homozygous) of the most strongly associated SNPs. In the genotype plot, each column represents a SNP and each row an individual, grouped by the presence or absence of the white submarginal band. **A)** Central America. **B)** Cauca Valley. **C)** Magdalena Valley and Venezuela.

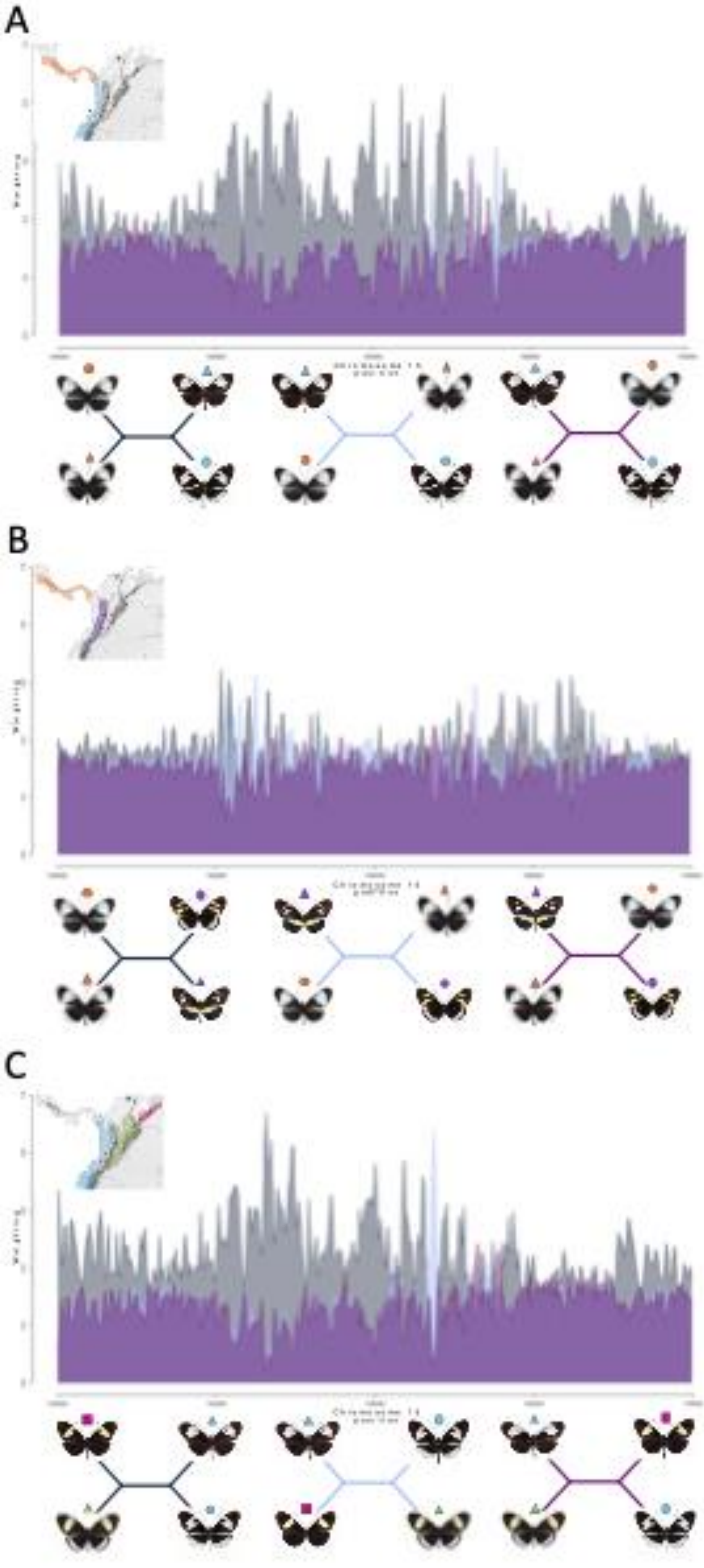

**Supplementary Figure 10. Twisst analysis results by geographic region.** The upper panel shows the geographic regions evaluated and topology weights across the candidate region (Hmel215003o: 1,500,000–1,600,000). The lower panel compares three hypotheses: clustering by geographic origin (dark blue), phenotype (light blue), and the control hypothesis (purple). **A)** Central America and Pacific. **B)** Central America and Cauca Valley. **C)** Magdalena Valley and Venezuela.

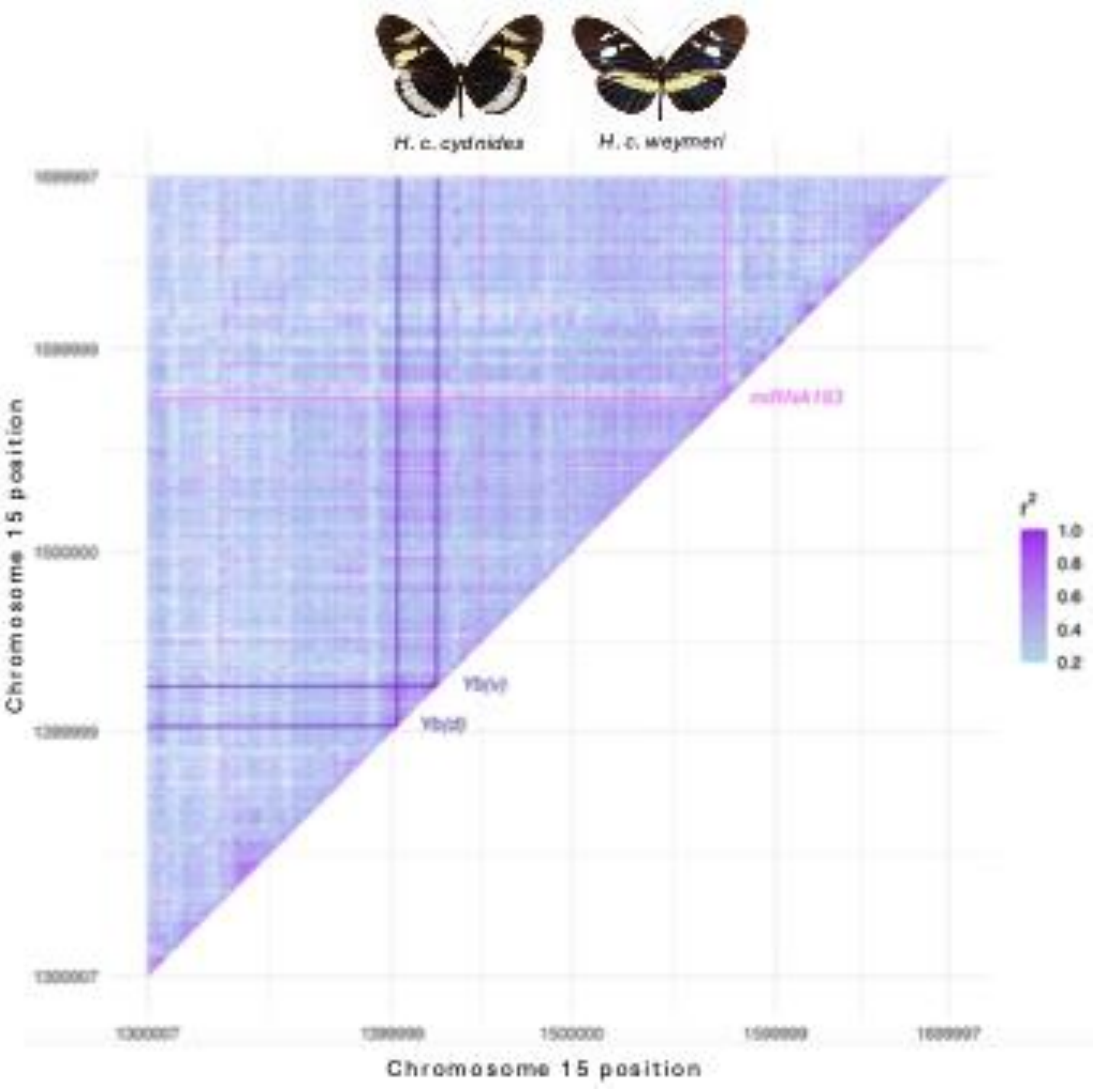


**Supplementary Figure 11. Linkage disequilibrium (r2) across the *ivory:mir-193* locus (Hmel215003o: 1,500,000–1,600,000).** The upper panel shows the subspecies evaluated and their corresponding phenotypes, while the lower panel presents the LD heatmap, indicating the positions of the regulatory elements Yb(d) and Yb(v), associated with the yellow bar, and the miRNA193-associated region, linked to the white submarginal band.

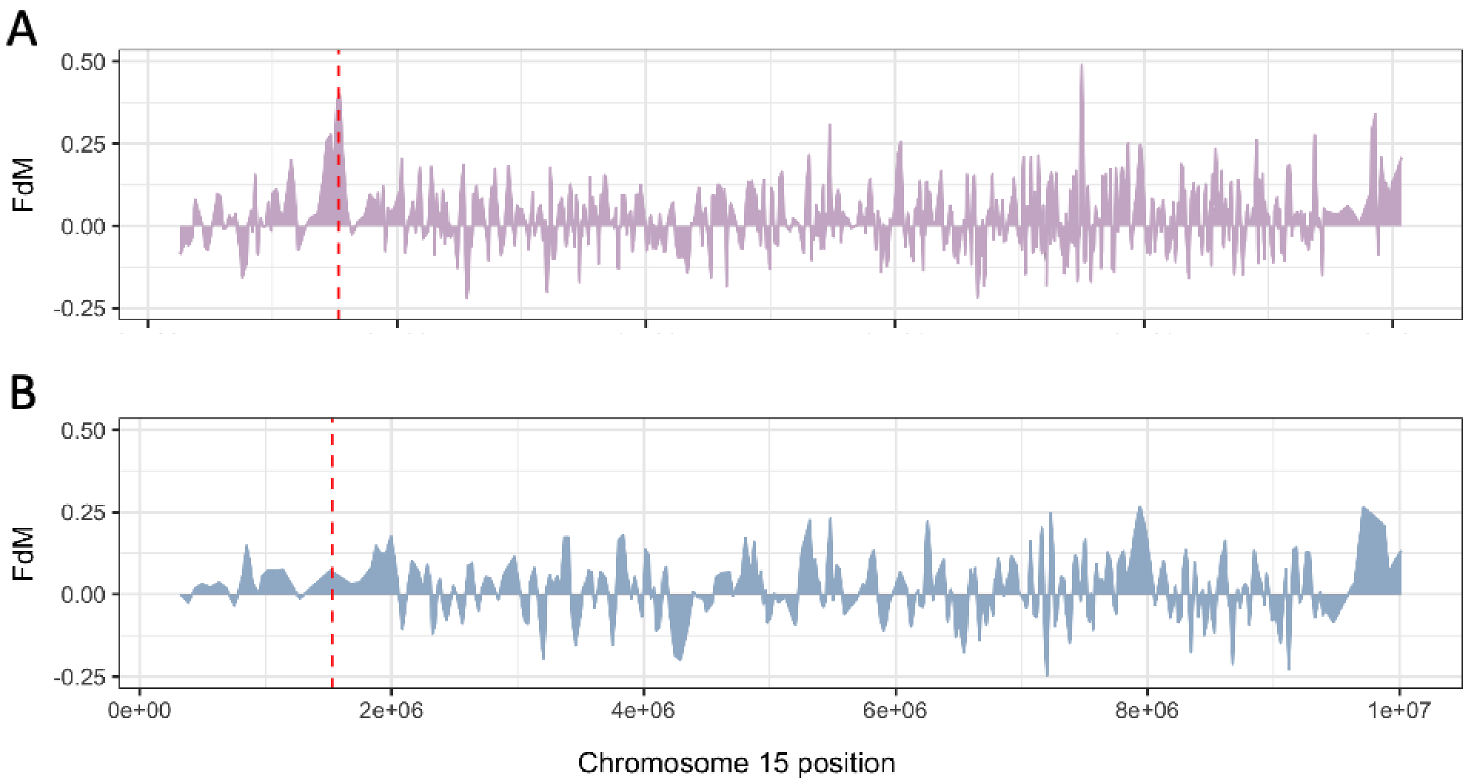


**Supplementary Figure 12. FdM statistic values across chromosome 15**. **A)** Analysis of *H. c. cydnides*, which exhibits the white submarginal band **B)** Analysis of *H. c. weymeri*, which lacks this trait. The dashed line marks the position of the highest peak, located at the *ivory:mir-193* locus. The purple line represents the results for *H. c. cydnides*, which exhibits the white submarginal band, while the blue line represents the results for H. *c. weymeri*, which lacks this trait. See Supplementary Figure 2 for the subspecies evaluated.

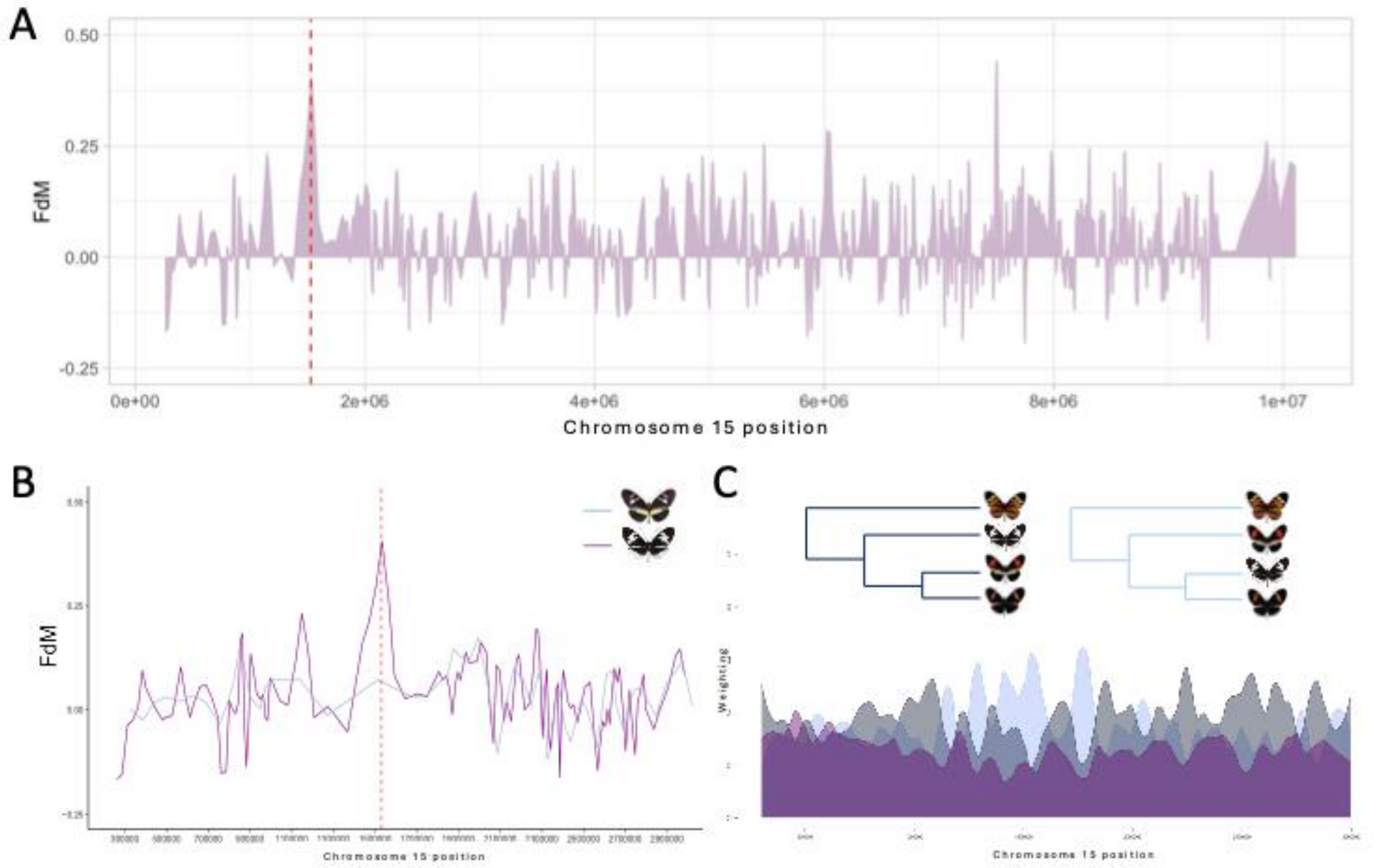


**Supplementary Figure 13. Introgression signals between *H. c. alithea* and *H. m. cythera*. A)** FdM statistic values across chromosome 15. The dashed line marks the position of the highest peak, located at the *ivory:mir-193* locus. **B)** FdM statistic values across the candidate region. The purple line represents the analysis of *H. c. alithea*, which exhibits the white submarginal band, while the blue line represents the analysis of *H. c. weymeri*, which lacks this trait. See Supplementary Figure 2 for the subspecies evaluated. **C.** Twisst analysis results for the same region and the hypothesized topologies evaluated.
